# Anatomical and neurophysiological data on primary visual cortex suffice for reproducing brain-like robust multiplexing of visual function

**DOI:** 10.1101/2021.12.07.471653

**Authors:** Guozhang Chen, Franz Scherr, Wolfgang Maass

## Abstract

Neural networks of the brain that process visual information have structural properties that differ significantly from those of neural networks which are commonly used for visual processing in AI, such as Convolutional Neural Networks (CNNs). But it has remained unknown how these structural differences are related to network function. We analyze visual processing capabilities of a large-scale model for area V1 that arguably provides the most comprehensive accumulation of anatomical and neurophysiological data that is currently available. Its network structure turns out to induce a number of characteristic visual processing capabilities of the brain, in particular the capability to multiplex different visual processing tasks, also on temporally dispersed visual information, with remarkable robustness to noise. This V1 model also exhibits a number of characteristic neural coding properties of the brain, which provide explanations for its superior noise robustness. Since visual processing in the brain is substantially more energy-efficient than implementations of CNNs in common computer hardware, such brain-like neural network models are likely to have also an impact on technology: As blueprints for visual processing in more energy-efficient neuromorphic hardware.

**Teaser:** A new generation of neural network models based on neurophysiological data can achieve robust multiplexing capabilities.

## 1 Introduction

The comprehensive model (*1*) for a patch of cortical area V1 in mouse provides an unprecedented window into the dynamics of this brain area. We show that it also provides a unique tool for studying brain-style visual processing and neural coding. The architecture of V1 exhibits an interesting combination of feedforward and recurrent connectivity: Neurons are distributed over several parallel 2D sheets, commonly referred to as layers or laminae in neuroscience. The neurons are recurrently connected, but not randomly or in an all-to-all manner. Rather, synaptic connections exist primarily between nearby neurons, both within a layer and between layers. Connectivity between layers supports a strong feedforward stream of visual information from L4 to L2/3 to L5/6, that is complemented by a host of recurrent loops. The dominance of short connections makes it possible to combine in V1 extensive recurrent connectivity with a really small total wire length, which is essential for its physical realization.

The model of (*1*) integrates besides these anatomical details also a host of neurophysiological data about area V1. The point neuron version of this model that we are considering employs generalized leaky integrate-and-fire neurons, more precisely GLIF_3_ neurons. These have in addition to the membrane potential two further hidden variables that model slower processes in biological neurons. The large diversity of neurons in the brain is reflected in the model of Billeh et al. through the use of 111 different types of GLIF_3_ neuron models that have each been fitted to experimental data in the Allen Brain Atlas (*2*).

The original model of (*1*) is not able to solve nontrivial computing tasks, since its synaptic weights were chosen on the basis of sparse experimental data about the mean and variance of synaptic weights. In contrast, synaptic weights in the living brain are individually tuned through a host of synaptic plasticity processes, and these processes induce higher-order correlations between weights that are crucial for computing capabilities of the network. At present we do not have enough data about these plasticity processes to reproduce them in a model. But we can address the question of what visual processing capabilities are supported by the model if synaptic weights are aligned for visual processing tasks through stochastic gradient descent. We applied this strategy to 5 different visual processing tasks that have commonly been considered in biological experiments (*3, 4, 5, 6, 7, 8*). Afterward, our model achieved high accuracy simultaneously for all 5 tasks, while working in a biologically realistic sparse firing regime close to criticality (*9, 10*). Surprisingly, its performance level remained in the same high-performance regime as the brain, even when we subjected the V1 model to noise in the images and in the network that it had not encountered during training. We demonstrate that this out-of-distribution (OOD) generalization capability of the V1 model with regard to new perturbations is far superior to that of CNNs. We provide an explanation for that through an analysis of neural coding properties of these two types of models: Both use high-dimensional neural codes for images. But the neural representation in the model of Billeh et al. is more robust because it employs, like the brain (*11*), a power law for the explained variance in higher PCA components that is close to a theoretically optimal compromise between the opposing goals to create noise-robust neural codes and to capture many details of visual inputs. In contrast, neural codes in CNNs have a different power law which reveals a focus on the latter (*12*). In addition, we demonstrate that the model of Billeh et al. uses preferentially those dimensions of population activity for coding that are orthogonal to the largest noise dimensions, like the brain does (*3*).

Altogether our results show that the currently available anatomical and neurophysiological data, as compiled in (*1*), provide the basis for a new generation of neural network models for visual processing that can multiplex diverse visual processing capabilities in a highly robust manner. Furthermore, these neural network models provide new paradigms for neuromorphic computing since they combine multiplexing capability and robustness to noise with small total wire length and highly energy-efficient sparse activity.

## 2 Results

### 2.1 Integration of anatomical and neurophysiological data, as well as data on noise in the brain, into a neural network model of area V1

Several decades of intense research efforts have accumulated a large body of knowledge about the anatomy and neurophysiology of the visual cortex, especially for primary visual cortex, i.e., for area V1. But it has remained unknown to what extent this insight into the structure of V1 can be related to its function. We have examined this question for the case of the large-scale model for a patch of V1 in mouse from (*1*), which is arguably the most comprehensive integration of anatomical and neurophysiological data on area V1 that is currently available. We will refer to the point neuron version of this model, also in combination with our data-based noise model that we discuss below, as the Billeh et al. model.

The Billeh et al. model is a spatially structured model for a patch of V1 that consists of 51,978 neurons from four main classes: One class of excitatory neurons and three classes of inhibitory neurons (Fig. 1B) that are distributed over 5 horizontal layers of neurons, labeled as L1, L2/3, L4, L5, and L6. Synaptic connections between these neurons are generated from data-based connection probabilities. These are defined in terms of base connection probabilities (Fig. 1C) that depend on the class and layer of the pre- and postsynaptic neuron. These base connection probabilities are scaled for each concrete pair of neurons by an exponentially decaying function of the lateral distance between their somata (Fig. 1D). This distance-dependent scaling entails that the vast majority of synaptic connections are between nearby neurons, and hence that the total wire length is small. But it also impacts the specific style of computational processing in the V1 model: Information is not continuously spread out all over the network as in randomly connected recurrent neural networks, that are frequently used as models for neural networks of the brain. To transform images and movies into input currents to neurons in this V1 model, we employed the preprocessing module (LGN model) of (*1*), see Fig. 1A. It consists of 17,400 filters that model in a qualitative manner the responses of four classes of experimentally observed LGN neurons (sustained ON, sustained OFF, transient ON/OFF, and transient OFF/ON), that are further subdivided according to preferred temporal frequencies.

**Figure 1:**
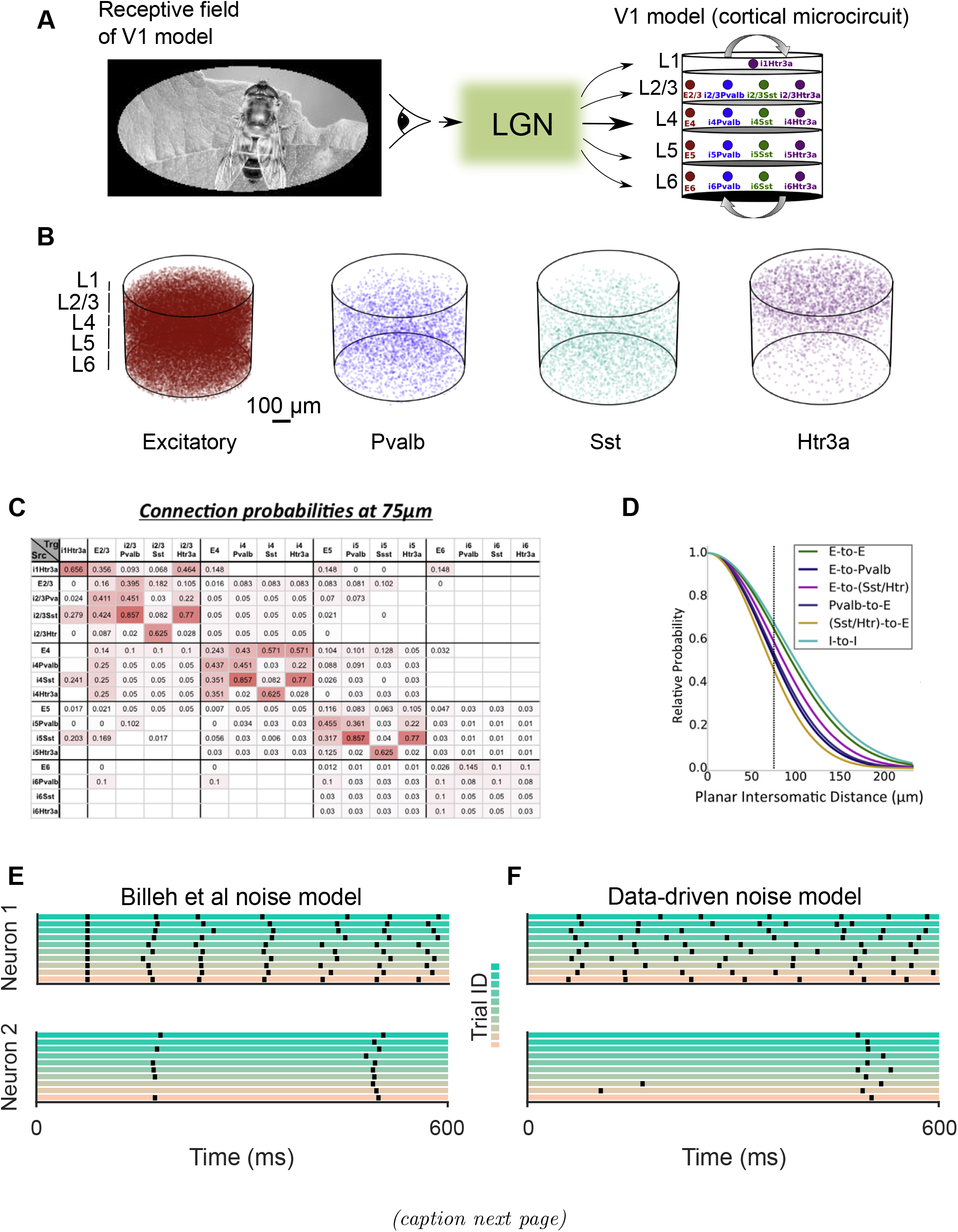
V1 model of (*1*). **(A)** The model consists of 4 classes of neurons on 5 layers. It comes together with a model for LGN, that transforms visual inputs into input currents to neurons in the V1 model. The LGN model receives visual input from an oval in the central part of an image (*1*). **(B)** The model contains one excitatory and three inhibitory neuron classes. Each dot denotes the position of a neuron. **(C)** The data-based base connection probabilities of (*1*) depend on the cell class to which the presynaptic (row labels) and postsynaptic neuron (column labels) belongs. White grid cells denote unknown values. **(D)** The base connection probability from **(C)** is multiplied according to (*1*)) for any given pair of neurons by an exponentially decaying factor that depends on the lateral distance between them. Panels A, C, D are reproduced from (*1*). **(E)** Spike outputs of 2 randomly selected neurons from the Billeh et al. model for 10 trials with the same input (a trial of visual change detection task for natural images), using the noise model of Billeh et al. **(F)** Same as in **(E)** but for the version of the data-based noise model with *s = q =* 2 that we used as default-noise model during testing. It causes substantially larger trial-to-trial variability.

Individual neurons are modeled as point neurons. But in contrast to the customary leaky integrate-and-fire (LIF) neuron models, the Billeh model employs 111 different variations of the LIF model, which are referred to as generalized leaky integrate-and-fire (GLIF_3_) neuron models because they have in addition to the membrane potential two other internal variables that model after-spike currents in the neuron on slower time scales. These 111 different neuron types have been fitted to experimental data for 111 selected neurons from the neocortex according to the cell database of the Allen Brain Atlas (*2*).

The neurons in the model of (*1*) received besides inputs from the LGN model and inputs from other neurons also a small noise current. This noise was generated by a single Poisson source for all neurons. Hence this noise is highly correlated, but its amplitude is so small that it has only little impact on neural firing (Fig. 1E). We used this noise model during training, but instead used a data-based noise model during testing. This noise model is based on experimental data from area V1 of the awake mouse (*11*). More precisely, we employed the heavy-tailed distribution of noise amplitudes that arises from their experimental data (Fig. S1). Furthermore, we superimposed two forms of noise: A quick form noise with scaling factor *q* where a new value is drawn every ms from this distribution, mimicking for example noise that arises from stochastic synaptic release, and a slow form of noise with scaling factor *s*, where a new value is drawn from this heavy-tailed distribution once at the beginning of each trial. The latter mimics the well-known dependence of neural responses to the state of the network at the beginning of a trial, see e.g. (*14*). We use the default values *s = q =* 2 for scaling these two forms of noise. The resulting noise model causes a qualitatively similar trial-to-trial variability of network responses in the V1 model as in the brain (compare Fig. S2B with Extended Data Fig. 5 of (*11*)). This trial-to-trial variability is shown for 2 sample neurons from the V1 model in Fig. 1F. The resulting Fano factor of spike counts in 10 ms windows has then a value of 1.46 in the V1 model, which is close to the measured value of 1.39 in mouse V1 (*15*). To get a clearer picture of the noise robustness of the V1 model, we tested its computational performance also for substantially larger values of the scaling factors *q* and *s* (Fig. 3E).

### 2.2 The V1 model can multiplex diverse computations on visual input streams

Classification of static images is a very popular test for neural networks. But brains have visual processing capabilities that go far beyond that, since visual information arrives in natural environments, especially in the presence of active vision, in a piecemeal manner. Hence brains need to be able to integrate temporally dispersed information, which can in general not be carried out by a feedforward neural network. Therefore we tested our model not only on a standard image classification task (handwritten digits from the MNIST dataset), but also on 4 tasks that require temporal integration of visual information. The latter ones have all been used in mouse experiments, and data on their behavioral performance are available. The 5 selected tasks are illustrated in Fig. 2: Discrimination of subtle differences in the orientation of drifting gratings (Fig. 2B), as in the experiments of (*3, 4*), a generic image classification task (Fig. 2C), visual change detection tasks for natural images and static gratings (Fig. 2D), as considered in (*5, 6*), and accumulation of temporally dispersed cues on the left and right (Fig. 2E), as considered in (*7, 8*), some of them with slightly longer delay periods.

**Figure 2:**
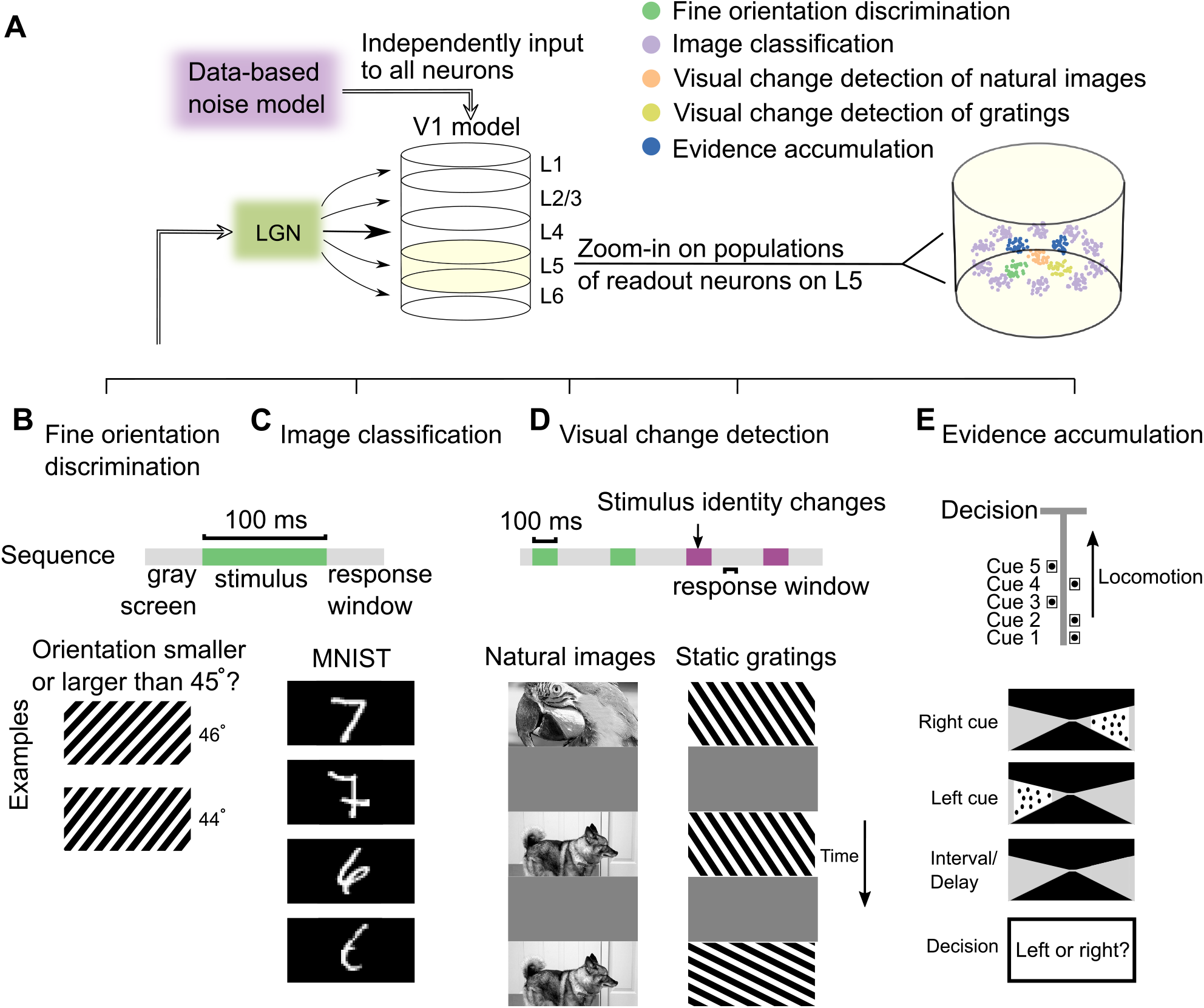
Illustration of the readout convention and the 5 visual processing tasks for which the V1 model was trained. **(A)** The visual stimuli for all 5 tasks were presented to the LGN model. Separate competing pools of pyramidal cells on L5 were chosen as readout neurons for each task. **(B-E)** Schematic diagrams and timings of 5 visual tasks (Materials and Methods). **(B)** In the fine orientation discrimination task, the network received a drifting grating with an orientation very close to 45°, and neurons in the corresponding readout pool had to fire if the orientation was larger than 45*°*. **(C)** For the image classification task, the network received a handwritten sample of a digit from 0 to 9 from the MNIST data set, and the corresponding one of the 10 readout pools for this task had to fire stronger than the others (two samples for digits 7 and 6 are shown). **(D)** For the visual change detection task, a long sequence of images was presented, with gray screens in between. A corresponding readout pool had to become active during the response window if the most recent image differed from the preceding one. Both natural images and static gratings were used. **(E)** In the evidence accumulation task, 7 cues were presented sequentially, and after a delay, a corresponding readout pool had to indicate through stronger firing whether the majority of the cues had been presented on the left or the right.

In order to test the performance of the V1 model on these tasks, one has to specify a convention for extracting the network decision. One frequently used convention, see e.g. (*16*), is to let an external “readout neuron” that receives synaptic input from all neurons in the network produce the network decision (Fig. S3C). Obviously, there are no such readout neurons in the brain that receive synaptic inputs from all neurons in a patch of the neocortex. Furthermore, this convention is not suitable for probing the computational capability of such a network model. Theoretical results ((*17*)) imply that if the network model is sufficiently large and has diverse units, such readout neurons tend to become computationally quite powerful when its weights have been properly trained, and is likely to mask the computational contribution of the neural network model itself. Therefore we demanded that in V1 model, like in the brain (*18*), projection neurons within the network extract computational results from the model, and project them to downstream networks. In particular, a large fraction of pyramidal cells on L5 projects to subcortical areas, and can therefore use the computational result of the network to trigger a behavioral response. Therefore we selected for each computational task and each possible outcome of a network decision a population of 30 excitatory neurons on L5 (Fig. 2A). If this population produced more spikes during the response window than competing populations that voted for other outcomes, then the outcome for which it “voted” was viewed as the network decision. One important difference to the convention of using global readout neurons is that the set of neurons in the network that provides synaptic inputs to a neuron within the network is substantially smaller and spatially constrained.

With the values of synaptic weights provided by (*1*), the V1 model is incapable of performing any of the 5 tasks; the accuracy is close to chance level. We then applied stochastic gradient descent, like in (*19*), to the synaptic weights of connections within the V1 model of Billeh et al., and to connections from the LGN model to the V1 model. No synaptic connections were added during this process. We also made sure that the signs of synaptic weights could not change, i.e., we maintained the validity of Dale’s law. We used a loss function for gradient descent that penalized inaccurate decisions by the chosen populations of readout neurons, see Materials and Methods. The loss function also penalized biologically unrealistic high firing rates. Stochastic gradient descent was implemented through a variation of BPTT, with the help of a suitable pseudo derivative for handling the discontinuous dynamics of spiking neurons, as suggested by (*20*). To avoid artifacts arising from gradient descent caused by the hard reset of GLIF_3_ neurons after a spike, we subtracted instead a fixed value from the membrane potential after each spike. Control experiments show that this modification causes no significant difference in the spike output of a neuron (Fig. S4). We trained the model of Billeh et al. with its original noise model (Fig. 1E), and tested its performance with the biologically more realistic noise model of Fig. 1F, and also in the presence of even more noise.

After training, the Billeh et al. model achieved on all 5 tasks a performance that was in the same range as reported behavioral data (Table 1), with an average accuracy of 89.10%. This performance did not depend on our particular choice of readout neurons in L5: Choosing randomly distributed instead of co-located pyramidal cells yielded an average accuracy of 91.56%. As expected, choosing instead global linear readout neurons for each task led to a substantially higher accuracy of 97.73%. The differences of readout scenarios can explain why behavioral performance lags behind neural coding fidelity in area V1 (*4*) (Supplementary Note 1). Sample computations of the V1 model for each of the 5 tasks are shown in Fig. S5-S9.

**Table 1:** The Billeh et al. model achieves high accuracy in all 5 tasks, consistent with the behavior performance of mice in similar tasks, after 6 training epochs.

|  | Test accuracy | Behavior accuracy | Mean firing rate (Hz) | Spike raster |
| --- | --- | --- | --- | --- |
| Fine orientation discrimination | 93.15% | $\sim 83\%^a$ | 3.97 | Fig. S5 |
| Image classification | 88.92% | N/A | 4.11 | Fig. S6 |
| Visual change detection of natural images | 84.13%/83.25%<sup>b</sup> | $\sim 73\%/77\%^c$ | 3.97 | Fig. S7 |
| Visual change detection of gratings | 89.25% | $\sim 60\%^d$ | 3.90 | Fig. S8 |
| Evidence accumulation | 90.92% | $\sim 85\%^e$ | 3.96 | Fig. S9 |

(a) Estimated from Fig. 4C of (33) when the orientation difference was $90^\circ$ .
(b) In the visual change detection task for natural images, the two values refer to testing with familiar and novel images, respectively.
(c) Estimated from Fig. 1I of (5) when familiar and novel images were presented to mice, respectively.
(d) Estimated from Fig. 3A of (34) when the orientation difference was $5^\circ$ .
(e) Estimated from Fig. 1C of (7) when the number of cues was 6.
Note that the behavioral experiments had longer time delays that made the tasks somewhat more difficult.

The median strength of inhibitory synapses increased from 0.03 to 1.65 pA during training; the median weight of excitatory synapses decreased from 2.96 to 1.43 pA, see Fig. 3A, and Fig. S10 for more detailed analyses in terms of the neuron types involved. Also, the distribution of neural firing activity was after training still close to the measured distribution in the brain, see Fig. 3B, with an average firing rate of 4 Hz.

**Figure 3:**
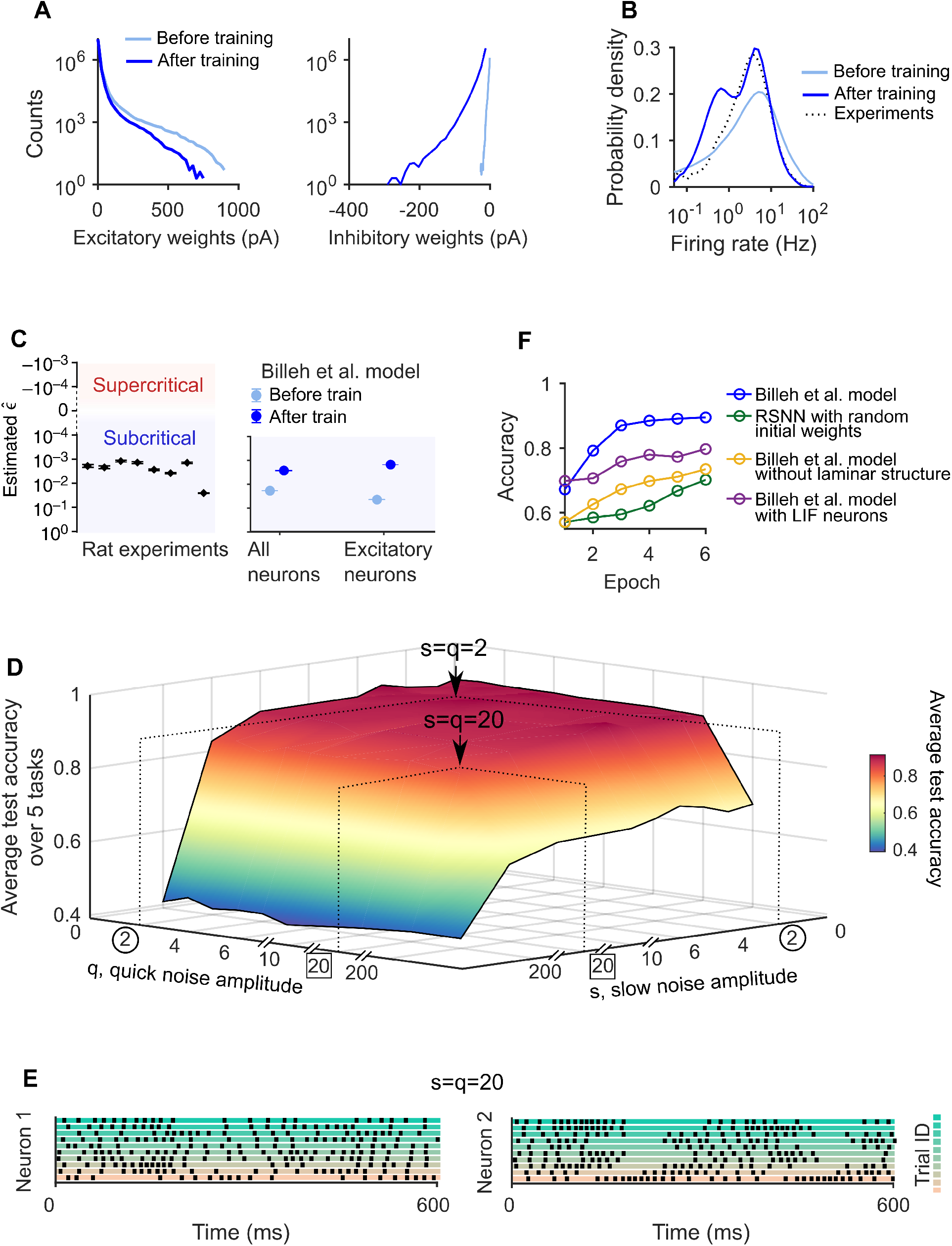
Analysis of the V1 model after training. **(A)** Distributions of excitatory weights (left) and inhibitory weights (right) before and after training. **(B)** Average distributions of firing rates for the 5 task before and after training. The distribution is moved through training closer to the one recorded in V1 (*6*). **(C)** Criticality of the Billeh model is analyzed and compared with experimental data. The y-axis shows estimates of 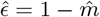, where 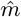 is the estimated branching ratio. Estimates of this value in the rat brain from (*13*) are reproduced on the left. The Billeh et al. model produces almost the same branching ratio as the brain, especially after training. Error bars on the left represent 16 to 84% confidence intervals. Error bars on the right represent SEM over 10 trials. **(D)** The average accuracy of the V1 model on test data for the 5 tasks, as function of the amplitude *q* of quick and the amplitude *s* of slow noise. The model had been trained just with the default noise of the Billeh model, which has much less impact on neural activity according to Fig. 1 E, F. The arrow in the back points to the accuracy for the default values *s = q =* 2 for the data-based noise model. One sees that the average accuracy for the 5 tasks is also robust to much larger noise amplitudes, e.g. for *s = q =* 20, see arrow in front, it still has an average accuracy of 83.07%. The resulting trial-to-trial variability of neural firing is substantial for this noise level, see **(E)** for samples of spiking activity of the same 2 neurons as in Fig. 1E, F for 10 trials with the same network input (image). **(F)** Average test accuracy across 5 tasks of the V1 model and of control models that lacked salient structural features of the V1 model; plotted as a function of the number of training epochs (Materials and Methods). Deleting salient structural features of the V1 model caused substantially slower progress of stochastic gradient descent training.

Experimental data suggest that neural networks of the brain typically operate in a critical regime (*21, 22, 10*). We evaluated the criticality of the V1 model by measuring its branching ratio of neural activity, as suggested by (*10*). We found that both the untrained and the trained V1 model operates in a slightly subcritical regime. Training moved the model somewhat closer to the critical regime, reaching values of the branching ratio that almost perfectly matched recorded data from the brain (Fig. 3C). Hence the V1 model operated in a dynamic regime that closely matches experimental data.

Interestingly, when the same training procedure was applied to control models that lacked salient structural features of the Billeh et al. model, their task performance advanced substantially slower, see Fig. 3F and Supplementary Note 2. This result suggests that the laminar cortical circuitry of V1 supports the efficiency of stochastic gradient descent. In particular, slow internal processes within GLIF_3_ neuron models contribute to it because gradients move more effectively through the corresponding slowly changing internal variables than through spikes. We propose that the dominance of short connections within and between laminae of laminar cortical microcircuits in the brain also supports gradient descent learning, since local errors in computation are not immediately and continuously spread out all over the network like in randomly connected networks without this characteristic architecture of cortical microcircuits.

We tested the robustness of the resulting multiplexed visual processing capability of the V1 model after training by exposing its neurons to substantially larger amplitudes *q* and *s* of the data-based noise model, although it had never been exposed to such noise during training (where we only applied the really small noise considered in (*1*)). Surprisingly, it is almost impossible to destroy its multiplexed visual processing capability (Fig. 3D): It remained stable even when the amplitudes *q* and *s* of quick and slow noise were increased by several orders of amplitude.

### 2.3 The power spectrum of neural codes provides an explanation for the astounding robustness of visual processing by the V1 model

An explanation for the robustness of visual processing in area V1 has been provided by (*11*). They verified through large-scale recordings from V1 in mouse a theoretically predicted link between noise robustness of visual processing and neural codes for images. They found that V1 employs high-dimensional neural codes for images, but the power of higher PCA components decays sufficiently fast so that its neural codes remain noise-robust. More precisely, they introduced a cross-validated principal component analysis that provides unbiased estimates of the stimulus-related variance. They found that the amount of explained variance continues to increase as further PCA dimensions were included without saturating below the dimensionality (= size) *d* of the image ensemble. We applied exactly the same analysis to the trained model of Billeh et al., and found that the model exhibited the same coding property (Fig. 4A). It was also shown in (*11*) that the explained variance of the *n*th principal component of network representations of images follows a power-law *n*^−*α*^. The exponent *α* characterizes how fast the variance that is explained by higher PCA dimensions decays. Their theoretical analysis predicts that *α =* 1 + 2*/d* is the optimal value (Fig. 4F), since this value provides a theoretically optimal compromise between encoding too many details (leading to smaller values of *α*), and keeping the neural code robust to perturbations (leading to larger values of *α*). Other theoretical work (*23*) also predicts that a value of *α* close to 1 enhances under mild conditions downstream generalization performance. In-vivo recordings of (*11*) found that the value of *α* for primary visual cortex of mouse is actually close to this optimal value *α =* 1 + 2*/d*. This neural coding property of area V1 in the brain has gained additional interest through the contrasting result of (*12*). They found that feedforward CNNs, which are viewed to be substantially less noise robust than the brain, have in fact a smaller *α*, as predicted by the theory. Hence we wondered whether our more brain-like neural network model for visual processing would exhibit a value that is closer to the theoretical optimum. We applied for that purpose the same measurement procedure as (*11*) to the V1 model, for a set of 2,800 randomly drawn natural images. Figure 4B shows the eigenspectrum of PCA component for the model of Billeh et al. before and after training, and also the measured eigenspectrum of V1 responses from (*11*). One sees that the eigenspectrum of the model is already before training quite close to that of the brain, and is moved by training even closer. The resulting exponent *α* of the power law (Fig. 4C) is for the model somewhat higher than in the brain. Fig. 4F, G, H suggest that this is largely due to the contributions of inhibitory neurons. They are generally found to have less precise neural codes for sensory stimuli, and consistent with that, their eigenspectra decayed substantially faster in the V1 model.

**Figure 4:**
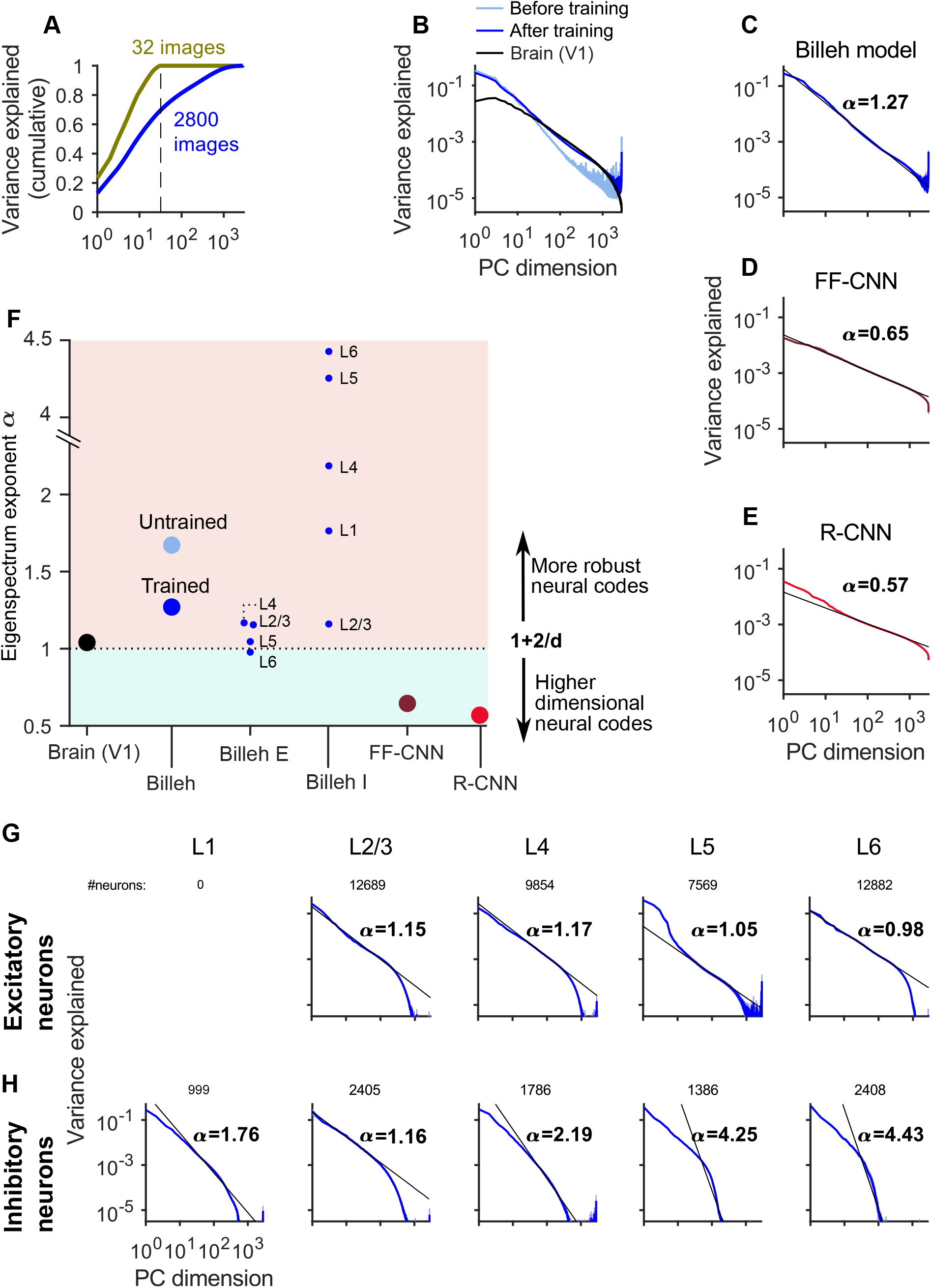
Comparing the eigenspectrum of neural codes in the V1 model of Billeh et al. with data from the brain, and with CNNs. **(A)** As in the brain, the cumulative fraction of explained variance saturates only at the dimension of the input ensemble, here shown for *d =* 32 and 2,800 natural images. **(B)** Eigenspectra of the untrained/trained V1 model of Billeh et al. with an ensemble of 2,800 randomly chosen natural images, and for mouse V1 (*11*). **(C, D, E)** Exponents of the power law for the V1 model of Billeh et al., for feedforward CNNs, and for recurrent CNNs; all for the same ensemble of 2,800 randomly drawn natural images. **(F)** Summary result for exponents of the power law for the brain the V1 model of Billeh et al., and for CNNs. **(G)** Eigenspectra of excitatory neurons on different layers of the model by Billeh et al. exhibiting values close to the measured value from a large sample of neurons in V1. **(H)** The same for inhibitory neurons. These exhibit on all layers except L2/3 substantially smaller coding fidelity.

**Figure 5:**
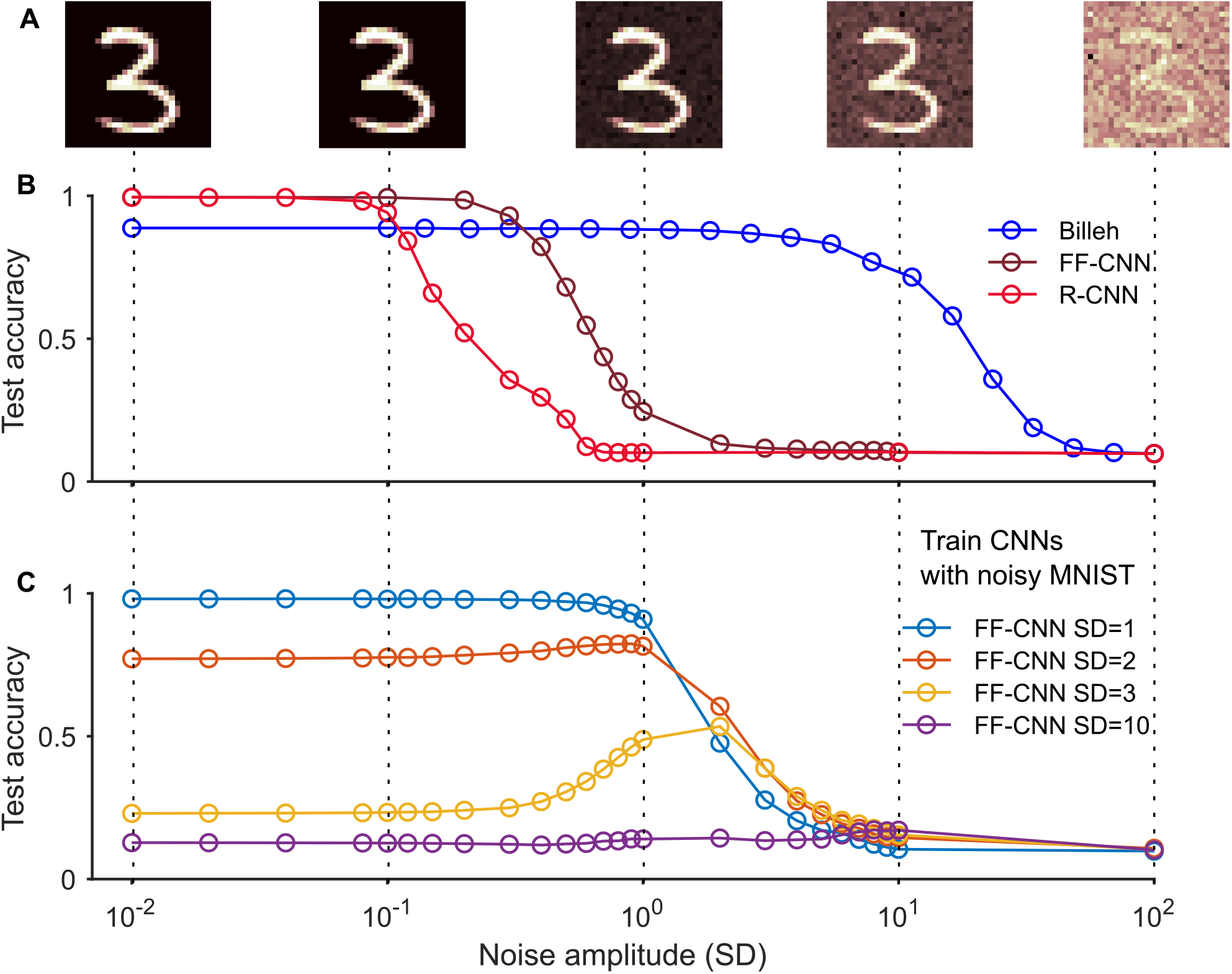
Robustness of the V1 model and of CNNs to noise in images. **(A)** Samples of handwritten MNIST digits with Gaussian noise drawn independently from 𝒩(0, SD) for each pixel, for different values of the SD. **(B)** While the V1 model never quite reaches the peak performance of the CNNs, it tolerates noise with fairly high SD, whereas the performance of feedforward (FF)-CNNs and recurrent (R)-CNNs is substantially degraded even by noise with small SD. **(C)** Even when CNNs are trained on images with a particular noise statistics (SD), they do not generalize well to test images with a different value of SD. Furthermore, they do not achieve for SD between 1 and 10 the same noise robustness as the V1 model even when they were trained on images with that type of noise.

We have also reproduced the result of (*12*) that feedforward CNN has a substantially smaller value of *α* (Fig. 4D) and found that the recurrent CNN model of (*24*), which also achieves very high accuracy for image classification, has an even smaller value of *α* (Fig. 4E). Altogether our results imply that the V1 model has, unlike CNNs, similar neural coding properties as area V1 in the brain. Furthermore, these can be linked according to the theory of (*11*) to its remarkable noise robustness.

### 2.4 Comparing noise robustness and OOD generalization of the V1 model and CNNs

Our preceding analyses of neural coding in the V1 model of Billeh et al. and CNNs suggest that the former is more noise-robust. Since it is hard to compare their robustness to noise within the networks with that of CNNs, because their computational units are so different, we compared instead their robustness to noise in the visual input, concretely to Gaussian pixel noise that was added to handwritten digits from the MNIST dataset. We used Gaussian noise with mean 0 and different SD. We first trained each type of neural network on the original dataset without noise, and then tested their classification performance on images with noise, see Fig. 5A for samples. Fig. 5B shows that the classification performance of the V1 model is substantially more robust to the added noise during testing, as predicted by the preceding analysis of the different neural coding strategies of the V1 model and CNNs.

Since neither of these networks had been trained with the noisy images for which they were tested, we are analyzing here a particular OOD (out-of-distribution) generalization capability of the V1 model and of CNNs. Fig. 5C demonstrates that even if CNNs are trained for a particular noise statistics, they do not perform well if they are tested on images with a different SD of Gaussian noise. In contrast, the V1 model exhibited perfect OOD generalization in this respect. Also, the remarkable robustness of the V1 model to noise within the network (Fig. 3D), which had not been present during training, can be viewed as an OOD generalization capability.

### 2.5 Neural coding dimensions for visual inputs are in the trained V1 model largely orthogonal to noise, like in the brain

A further explanation for robust coding capabilities of the brain was provided by experimental data of (*3*). They reported that dimensions in which differences between visual stimuli were encoded in the brain were nearly orthogonal to the largest noise mode, which therefore had little effect on coding fidelity. We carried out the same analysis for the V1 model, using the same visual stimuli (moving gratings with orientation differences that were close to the perception threshold). The definition of the discriminability index *d*′, according to (*3*) a proposed measure for the fidelity of neural population coding, is illustrated in Fig. 6A. Neural population responses **r**_A_(*t*) and **r**_B_(*t*) to two stimuli A and B form two distributions (ellipses). PLS (partial least square) analysis projects them onto a subspace where they become most distinct. The discriminability *d*′ is defined as the separation, Δ***µ***, of the two distributions along the dimension orthogonal to the optimal boundary (green line) for classifying stimuli in this subspace, divided by the SD of each distribution along this dimension.

**Figure 6:**
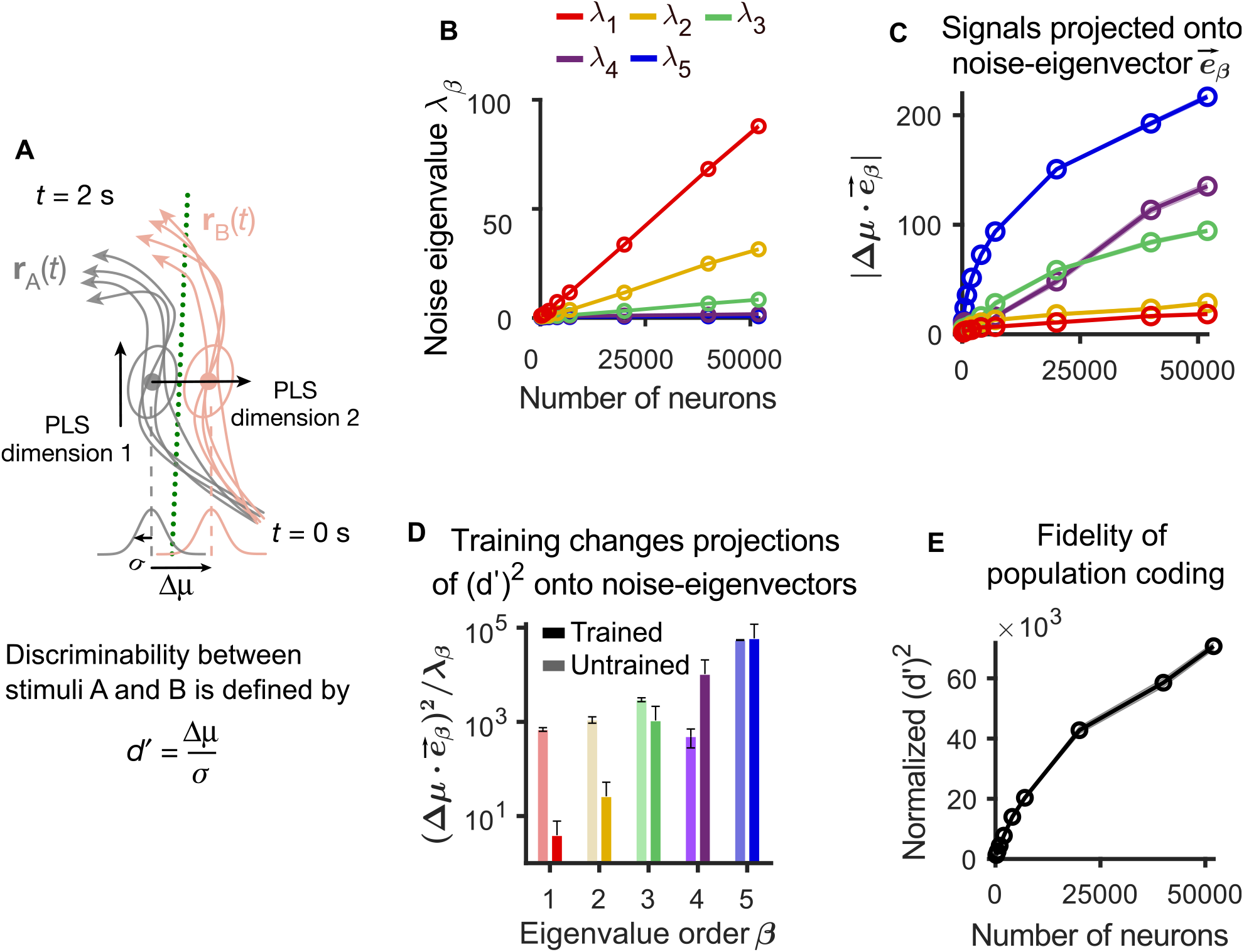
Analyses of relations between signaling and noise dimensions in the V1 model, and of the impact of noise correlations when one considers larger numbers of neurons. **(A)** Schematic of the calculation of the discriminability index *d*′ according to (*3*). **(B)** The 5 largest eigenvalues *λ*^*β*^ (*β =* 1, 2, …, 5) of the noise covariance matrix in the trained V1 model increase linearly with the number of sampled neurons. **(C)** Neural coding dimensions (Δ***µ***) in the trained V1 model were nearly orthogonal to the dominant eigenvectors of the noise covariance matrix, 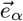, invariant to neuron numbers. **(D)** Training moved the neural coding dimensions so that they became more orthogonal to the dominant noise dimensions. **(E)** The squared discriminability index (*d*′)^2^ kept increasing when applied to more neurons om the V1 model. *d*′ values were normalized by those obtained for trial-shuffled data (averaged across 1 s). Shaded areas in **(B, D, E)** and error bars in **(C)** represent the standard error of mean (SEM) over 100 trials.

The eigenvalues of the noise covariance matrix are plotted in Fig. 6B as function of the number of neurons that are sampled in the V1 model. We found that also in the V1 model the projection of the signal difference Δ***µ*** onto the eigenvectors for the largest noise eigenvalues is relatively small (Fig. 6C). Furthermore, compared with the untrained Billeh model, training of this model moved the signaling dimensions to become more orthogonal to the largest noise dimension, see Fig. 6D. The projection of (*d*′)^2^ on an eigenvector can be interpreted as the signal-to-noise ratio because it is the ratio of the projected signal difference and noise eigenvalue (Materials and Methods).

It had been argued in (*3*) that the amount of visual information that is encoded in area V1 reaches a ceiling due to noise correlations. This result appeared to contradict the results of (*11*). A possible explanation of this discrepancy was offered in (*4*): They conjectured that the seemingly limited coding capability of V1 could be explained by the relatively small number of up to 1,300 neurons from which simultaneous recordings had been carried out in (*3*). They suggested that the apparent “ceiling” would rise if one records from more neurons. We tested this hypothesis in the V1 model of Billeh et al., and found that indeed the corresponding measurement *d*′ for the total amount of encoded information keeps increasing, although at a somewhat slower rate, when the number of neurons from which one records in the model rises from 1,300 to 51,978, see Fig. 6E.

## 3 Discussion

We have demonstrated that the neural network model of (*1*), which arguably provides the largest currently available accumulation of anatomical and neurophysiological data on area V1 in the mouse brain, provides not only a window into brain dynamics, but also into visual processing capabilities and neural coding properties that are entailed by these data. CNNs are currently in AI one of the most commonly considered types of neural networks for visual processing. They were also inspired by some aspects of visual processing in the brain, especially the existence of simple and complex cells in area V1. But a closer look shows that they differ from V1 in the brain in almost all other respects: With regard to their computational units (artificial neurons in CNNs versus spiking neurons in the brain), the diversity of their units (very few simple units versus a large diversity of neurons with different temporal dynamics), their large-scale architecture (usually feedforward, versus recurrent with laminar structure), their small-scale architecture (very simple network motifs versus a complex combination of feedforward and recurrent processing in cortical microcircuits), and total wire length (almost quadratic versus just linear growth with the number of processing neurons). But V1 in the brain also differs from CNNs with regard to two important visual processing capabilities: The brain can multiplex strategies for solving diverse visual tasks within the same network, in particular also tasks that require integration of sequentially arriving visual information. In addition, visual processing in the brain is very noise-robust, also to new types of noise (OOD generalization). We have shown here that the previously listed fundamental differences between the structure of V1 in the brain and CNNs are causally related to these two superior visual processing capabilities of the brain: The V1 model of (*1*), that integrates these structural features of V1 in the brain, is able to multiplex robust visual processing.

Since we can reproduce now these two important functional capabilities of area V1 in a model, we have a new research platform at our disposal for studying how neural coding properties of the brain emerge from its anatomical and neurophysiological features, and how they are related to its visual processing capabilities. We have demonstrated here the feasibility of this new research strategy by applying to the V1 model, an analysis of its neural coding that had already been used successfully for elucidating neural codes for images in area V1 of the brain: We analyzed the eigenspectrum of the explained variance of principal components of its neural codes for images. We found that the listed structural features of V1 in the brain do in fact induce a salient feature of its neural coding strategy: The V1 model exhibits a similar power law for neural codes of images as the brain. In contrast, CNNs have a power law with a substantially slower decay of the eigenspectrum. According to the theoretical analysis of (*11*), this implies that neural codes in CNNs are less noise-robust. Our V1 model also demonstrates that a further salient aspect of neural coding in area V1 of the brain results from its anatomical and neurophysiological details: According to (*3*) about 90% of the noise fluctuations in area V1 of the brain are constrained to dimensions of the population activity that are orthogonal to noise dimensions. We found that this is in fact an emergent property of the V1 model of (*1*).

On a more general level, we have shown that the V1 model of (*1*) can be seen as the first prototype of a new generation of neural network models for visual processing that capture substantially more features of brain processing than CNNs. Further work is needed to tease apart the functional implications of each of its structural features, and to port similar advanced brain-like visual processing capabilities into simpler neural network models. Often one uses instead of data-based models for neural networks of the brain randomly connected recurrent networks of strongly simplified neuron models. In our experience, neural coding and computational properties of recurrent neural networks vary substantially in dependence of their connectivity structure and neuron models. This highlights the need to test brain-like features not only in abstract models, but also in neural network models that integrate our available knowledge about the actual structure of these neural networks in the brain.

We propose that our method can also be applied to investigate how anatomical and neurophysiological details of interconnected higher and lower brain areas carry out distributed computations, in particular how higher cortical areas enhance visual processing capabilities of V1. A substantial body of anatomical and neurophysiological data on higher cortical areas and their connectivity to V1 is currently available for that, see e.g. (*6, 25, 26*). We expect that deficits in visual processing capabilities of the V1 model, such as limited spatial integration of image features and a relatively short working memory time span, will disappear when the V1 model is combined with models of higher brain areas.

The analysis of neural coding in the V1 model has produced a number of predictions for future biological experiments. In particular, we have shown in Fig. 6E that correlated noise reduces the coding fidelity of the network, but does not produce an a-priori bound for its sensory discrimination capability (this had already been hypothesized by (*4*)). A further prediction of our model is that the PCA eigenspectrum of neural codes for inhibitory neurons does not obey a power law for higher dimensions, see Fig. 4H. Finally, the values of excitatory synaptic weights in V1 are predicted to generally shrink through training (Fig. 3A left), while inhibitory weights are predicted to become stronger (Fig. 3A right).

Visual processing in the brain exhibits also with regard to two aspects of physical implementations substantial advantages over CNNs: Most synaptic connections in V1 are between nearby units, which is essential for an efficient physical realization of synaptic connections in neuromorphic hardware. This architectural feature is also likely to support faster learning (Fig. 3F). In addition, computations are carried out in the V1 model through event-based processing with very sparse firing activity. This computing regime is not only very energy efficient, but it also supports computations on tasks where temporal aspects play an important role, because it allows to let time represent itself in network computations. Hence, our analysis of a neural network model for area V1 of the brain paves the way for substantially more energy-efficient neuromorphic implementations of visual processing (*27*).

## 4 Materials and Methods

### 4.1 Neuron models

We based our study on the “core” part of the point-neuron version of the realistic V1 model introduced by (*1*). To make it gradient-friendly, we replaced the hard reset of membrane potential after a spike emerges with the reduction of membrane potential *z*_*j*_(*t*)*v*^*th*^, where *z*_*j*_(*t*) = 1 when neuron *j* fires at time *t* and *z*_*j*_(*t*) = 0 otherwise. *v*^*th*^ is the firing threshold of membrane potential. This causes no significant change in the neural response (Fig. S4). We simulated each trial for 600 ms. The dynamics of the modified GLIF_3_ model was defined as

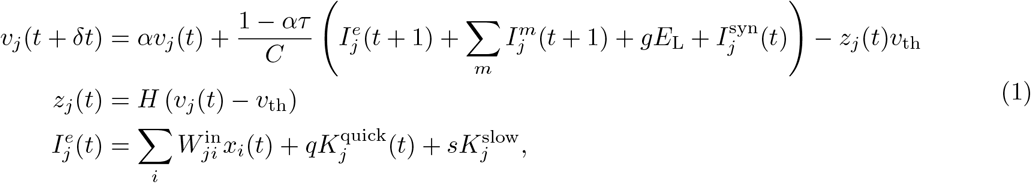

where *C* represents the neuron capacitance, *E*_*L*_ the resting membrane potential, *I*^*e*^ the external current, *I*^syn^ the synaptic current, *g* the membrane conductance, and *v*^th^ the spiking threshold. 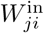 is the synaptic weight from LGN neuron *i* to V1 neuron *j*. The scales of the quick noise 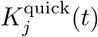 and the slow noise 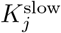 to neuron *j* are *q =* 2 and *s =* 2, respectively, unless otherwise stated. *K*_*j*_ was randomly drawn from the empirical noise distribution which will be elaborated on later. The decay factor *α* is given by *e*^−*δt/τ*^, where *τ* is the membrane time constant. *δt* denotes the discrete-time step size, which is set to 1 ms in our simulations. *H* denotes the Heaviside step function. To introduce a simple model of neuronal refractoriness, we further assumed that *z*_*j*_(*t*) is fixed to 0 after each spike of neuron *j* for a short refractory period depending on the neuron type. The after-spike current *I*^m^(*t*) was modeled as

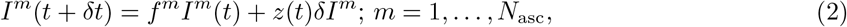

where the multiplicative constant *f*^m^ = exp (−*k*^m^*δt*) and an additive constant, *δI*^m^. In our study, *m =* 1 or 2. Neuron parameters have been fitted to experimental data from 111 selected neurons according to the cell database of the Allen Brain Atlas (*2*), see (*28, 1*), including neuron capacity *C*, conductance *g*, resting potential *E*_L_, the length of the refractory period, as well as amplitudes *δI*^m^ and decay time constants *k*^m^ of two types of after-spike currents, *m =* 1, 2.

### 4.2 Synaptic inputs

The Billeh et al. model specifies the connection probability between neurons, based on experimental data. The base connection probability for any pair of neurons from the 17 cell classes is provided in (*1*) by a table (reproduced in Fig. 1C); white grid cells denote unknown values. The entries in this table are based on measured frequencies of synaptic connections for neurons at maximal 75 µm horizontal inter-somatic distance. This base connection probability was scaled by an exponentially decaying factor in terms of the horizontal distance of the somata of the two neurons (Fig. 1D). This distance-dependent scaling is also based on statistical data from experiments (leaving aside finer details of connection probabilities). The synaptic delay was spread in [1, 4] ms, which was extracted from the Fig. 4E of (*1*) and converted to integers as the integration step is 1 ms.

The postsynaptic current of neuron *j* was defined by the following dynamics (*1*):

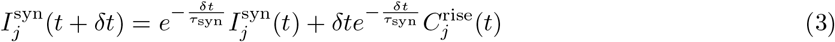

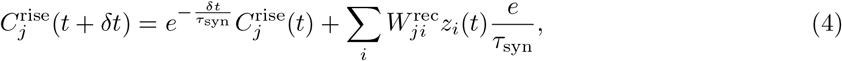

where *τ*^syn^ is the synaptic time constant, 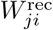 is the recurrent input connection weight from neuron *i* to *j*, and *z*_*i*_ is the spike of presynaptic neuron *i*. The *τ*_syn_ constants depend on neuron types of pre- and postsynaptic neurons (*1*).

### 4.3 Initial conditions

The initial conditions of spikes and membrane potentials were zero unless stated otherwise. The initial conditions of **W**^in^ and **W**^rec^ were given by the values in (*1*) unless stated otherwise.

### 4.4 Data-driven noise model

The noise currents 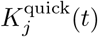 and 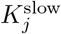 in Eq. 1 were randomly drawn from an empirical noise distribution. The quick noise 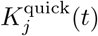 was drawn independently for all neurons in every 1 ms; the slow noise 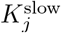 was drawn independently for all neurons once 600 ms. The empirical noise distribution (Fig. S1) was from the additive noise decoded from experimental data of mice response to 2,800 nature images (*11*) (https://figshare.com/articles/Recordings_of_ten_thousand_neurons_in_visual_cortex_in_response_to_2_800_natural_images/6845348). The decoding method was cross-validation principal component analysis (cvPCA) (*11*) which will be elaborated later. It measures the reliable variance of stimulus-related dimensions, excluding trial-to-trial variability from unrelated cognitive and/or behavioral variables or noise. We collected the variability (additive noise) to form the empirical noise distribution. We refer to the methods and supplementary materials of (*11*) for a detailed mathematical analysis of this method.

### 4.5 Readout populations

By default we employed 15 readout populations in the V1 model, whose firing activity during the response window encoded the network decisions for the 5 visual processing tasks. Each population consisted of 30 randomly selected excitatory neurons in layer 5, located within a sphere of a radius of 55 µm, with some distance between these spheres for different readout populations (Fig. S3A). The results were not sensitive to the number of neurons in each population (*19*). We also considered the case where the neurons in these readout populations were randomly distributed in L5 (Fig. S3B), and the case where each population was replaced by global linear readout neurons which received synaptic inputs from all neurons with activity (**Z**) in the V1 model, i.e. **Y**_global_ = **W**_readout_**Z** + **B, B** is the bias (Fig. S3C).

### 4.6 Visual processing tasks

We designed details of these 5 tasks to be as close as possible to corresponding biological experiments while keeping them as simple as possible. Only for the image classification task (MNIST) there exist no corresponding mouse experiments.

#### LGN model

The visual stimuli were preprocessed by the LGN model (Fig. 2A) according to (*1*) (it is actually meant to model preprocessing by the retina and LGN in a qualitative manner). This LGN model consists of 17,400 spatiotemporal filters that model responses of LGN neurons in mouse to visual stimuli (*29*). Each filter produces a positive output that is interpreted as firing rates of a corresponding LGN neurons.

According to the requirements of this LGN model, each visual input pixel was first converted to gray-scale and scaled into an interval [−*Int, Int*], *Int* > 0. The output of the LGN model was injected into the V1 model as external currents, i.e.,

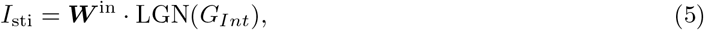

where *G*_*Int*_ represents images scaled into [−*Int, Int*] for *Int =* 2.

#### Fine orientation discrimination task

In mouse experiments, mice were trained to distinguish orientation of drifting grating stimuli (*3, 4*). The stimuli were presented for 750 ms or longer. To reproduce this task under the limitations of GPU memory, we input drifting grating to the V1 model through the LGN model for 100 ms (Fig. 2B). As in (*4*), stimuli were sinusoidal drifting gratings (spatial frequency, 0.05 cycles per degree, drifting rate, 2 Hz). Both in the training and testing processes, the orientation was uniformly drawn from [43, 47]*°* (i.e., 45 ± 2) with the precision of 0.1*°*. The orientation difference was the same as in (*4*). The initial phase was randomly sampled. The simulation sequence included 50-ms delay, 100-ms drifting gratings, and 50-ms response window in order.

In the response window, we defined the mean firing rate of readout population as

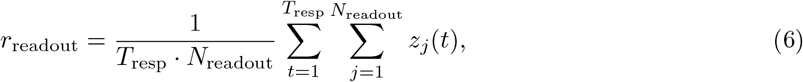

where the sum over *j* is over the *N*_readout_ = 30 readout neurons and the sum over *t* is over the time length of response window *T*_resp_ = 50 ms. If *r > r*_0_ = 0.01, then this reported a network decision that the orientation was larger than 45°. Otherwise, it reported that the orientation was smaller than 45°.

#### Image classification task

To demonstrate that the V1 model is also able to classify images, we included the task to classify handwritten digits from 0 to 9 from the MNIST dataset (Fig. 2C). The timing of input images and response windows was the same as in the preceding task. The task was to decide which digit was denoted by the handwritten image (two samples for 7 and 6 are shown in Fig. 2C). Each of the 10 readout populations for this task was assigned to one of the 10 digits. The network decision was taken to be that digit for which the readout population fired most strongly during the response window.

#### Visual change detection task with natural images

In mouse experiments (*30, 6*), mice were trained to perform the visual change detection task with natural images. A sequence of static natural images (250 ms), interleaved by short phases (500 ms) of gray screens, was presented as visual input; mice had to report whether the most recently presented image was the same as the previously presented one. To reproduce this task under the limitations of GPU memory, we presented natural images for 100 ms each, with the gray delays between them lasting for 200 ms (Fig. 2D). Note that the first image was presented after 50 ms. All images were selected from a set of 40 randomly chosen images from the ImageNet dataset (*31*). The probability that the next image differed from the preceding one was set to 50%. In case of a changed image identity, the model had to report within a time window of 50 ms length that started 150 ms after image onset (response window). If the mean firing rate of the readout population in the response window *r*_readout_ *> r*_0_, it reported a network decision that the image had changed. The computation of the V1 model on this task has been further analyzed in (*19*).

#### Visual change detection task with drifting gratings

We also replaced the natural images above with static gratings which have different orientations and kept the input sequence the same (Fig. 2D). The setting of the static grating is the same as in the fine orientation discrimination task except it is static. The changing probability of orientation is 50%; the orientation of static gratings was uniformly drawn in [120, 150] (i.e., 135 ± 15) with the precision of 0.1*°*.

#### Evidence accumulation task

A hallmark of cognitive computations in the brain is the capability to go beyond a purely reactive mode: to integrate diverse sensory cues over time, and to wait until the right moment arrives for an action. A large number of experiments in neuroscience analyze neural coding after learning such tasks (see e.g., (*7, 8*)). We considered the same task that was studied in the experiments of (*7, 8*). There a rodent moved along a linear track in a virtual environment, where it encountered several visual cues on the left and right (Fig. 2E). Later, when it arrived at a T-junction, it had to decide whether to turn left or right. The network should report the direction from which it had previously received the majority of visual cues. To reproduce this task under the limitations of a GPU implementation, we used a shorter duration of 600 ms for each trial. The right (left) cue was represented by 50 ms of cue image in which the black dots on the right (left) side of the maze. Visual cues were separated by 10 ms, represented by the gray wall of the maze. After a delay of 250 ms, the network had to decide whether more cues had been presented on the left or right, using two readout populations for left and right. The decision was indicated by the more vigorously firing readout pool (left or right) within the response window of 50 ms.

### 4.7 Loss function

The loss function was defined as

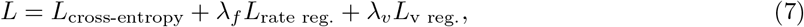

where *L*_cross-entropy_ represents the cross-entropy loss, *λ*_*f*_ and *λ*_*v*_ represent the weights of firing-rate regularization *L*_rate reg._ and voltage regularization *L*_v reg._, respectively. As an example, the cross-entropy loss of visual change detection tasks was given by

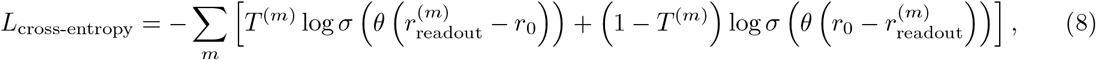

where the sum over *m* is organized into chunks of 50 ms and 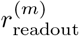 denotes the mean readout population firing rate defined in Eq. 6. Similarly, *T* ^(*m*)^ denotes the target output in time window *m*, being 1 if a change in image identity should be reported and otherwise 0. The baseline firing rate *r*_0_ was 0.01. *Σ* represents the sigmoid function. *θ* is a trainable scale (*θ > 0*) of firing rate.

We also used regularization terms to penalize unrealistic firing rates as well as unrealistic membrane voltages. Their weights, *λ*_f_ = 0.1 and *λ*_v_ = 10^*−*5^. The rate regularization is given by the Huber loss (*32*) between the target firing rates, *y*, calculated from the model in (*1*), and the firing rates, *r*, sampled the same number of neurons from the network model:

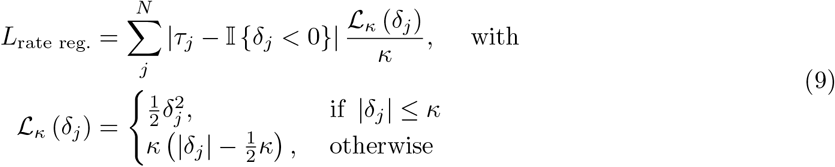

where *j* represents neuron *j, N* the number of neurons, *τ*_*j*_ *= j/N, δ =* 0.002, 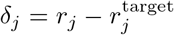. 𝕀 (*x*) = 1 when *x* is true; 𝕀 (*x*) = 0 when *x* is false.

The voltage regularization was given by

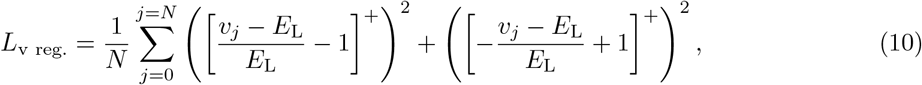

where *N* represents the total number of neurons, *v*_*j*_, the membrane potential of neuron *j, E*_L_, the resting membrane potential, [· · ·]^+^, rectifier function.

### 4.8 Training and testing

We trained the model for all 5 tasks together. Pairs of visual inputs and target outputs were collected in separate 64 batches for each task and these batches were interlaced during training. Apart from the change detection tasks, the spikes and membrane potentials were reset to 0 after each trial that consisted of 600 ms.

We applied back-propagation through time (BPTT) (*19*) to minimize the loss function. The non-existing derivative 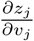 was replaced in simulations by a simple nonlinear function of the membrane potential that is called the pseudo-derivative. Outside of the refractory period, we chose a pseudo-derivative of the form

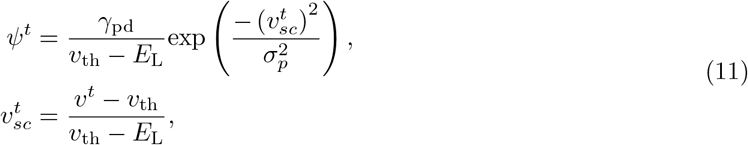

where the dampening factor *γ*_pd_ = 0.5, the Gaussian kernel width *σ*_*p*_ *=* 0.28. During the refractory period, the pseudo derivative was set to 0. During the training, we added the sign constraint on the weights of the neural network to keep Dale’s law. Specifically, if an excitatory weight was updated to a negative value, it would be set to 0; vice versa.

We would like to emphasize that we tested the trained model -whenever possible- for new visual stimuli that had not been shown during training (this was not possible for the gratings because there were not sufficiently many different visual stimuli for them). In that sense, we evaluated the generalization capability of the trained Billeh et al. model, rather than its capability to handle a fixed set of stimuli correctly (which often suffices to solve behavioral tasks in experiments). The model achieved on all 5 tasks a performance that is in the same range as reported behavioral data from corresponding mouse experiments (Table 1).

### 4.9 Other simulation details

The BPTT training algorithm was coded, as the simulation of the model, in TensorFlow, which runs very efficiently on GPUs, and also on multiple GPUs for training in parallel. We employed independent simulations in parallel by distributing trials for all 5 tasks over batches. Every batch consisted of 320 trials, 64 for each of the 5 tasks. In every trial, the model of Billeh et al. was simulated for 600 ms of biological time, which took, together with the calculation of gradients around 5 s on a NVIDIA A100 GPU. Once all batches had finished (one step), gradients were calculated and averaged to update the weights by BPTT. We define an epoch as 781 iterations/steps, because this represents one cycle through the full training dataset of MNIST. This computation had to be iterated for 6 epochs until the average performance on the 5 tasks was saturated. This took 20 h of wall clock time on 160 GPUs.

### 4.10 Control models

We used three control models in Fig. 3F. The first one was a randomly connected recurrent network of spiking neurons with the same numbers of neurons and connections, referred to as RSNN. In the RSNN, all data-based features of Billeh et al. model except for the number of neurons and synapses were removed: The GLIF_3_ neuron model was replaced by the standard LIF model; neural connectivity representing laminar structure with primarily local connection was replaced by random connectivity; diverse neuron types were replaced by a single neuron type: the excitatory neuron on L2/3 (node type id in Allen brain atlas: 487661754); Dale’s law was removed; initial weights were replaced by random values drawn from the Gaussian distribution with the same mean and variance as the initial values of the Billeh et al. model. In the second control model (Billeh et al. model without laminar structure), we kept the diverse GLIF_3_ neuron models, but the laminar connectivity structure was replaced by random connections, and we kept the same number of synaptic connections. In the third control model (Billeh et al. model with LIF neurons), the diverse GLIF_3_ neuron models were replaced by the standard LIF neuron model, all other features were kept.

### 4.11 Branching ratio as a measure for criticality

Based on the work of (*35*), where the branching ratio was recommended as a rather reliable measure for criticality of a network, we examined this branching ratio for the V1 model. In particular, this measure was shown there to be more robust to subsampling. The branching ratio is defined as the ratio of the number of neurons spiking at time *t* + 1 to the number of spiking neurons at time *t*. Critical regimes, by their nature, are balanced and avoid runaway gain (positive or negative) and have a branching ratio of 1.0. We stimulated the V1 model as in the visual change detection task of nature images for 15 s.

In a network with *A* active neurons at time *t*, if the branching ratio has a fixed value *m* then ⟨*A*_*t*+1_ | *A*_*t*_⟩ = *mA*_*t*_+*h* where <|> denotes the conditional expectation, *m* is the branching ratio and *h* is a mean rate of an external drive/stimulus. Considering subsampling, *a*_*t*_ is proportional to *A*_*t*_ *o*n average < *a*_*t*_ |*A*_*t*_⟩ = *ηA*_*t*_+*ξ*, where *η* and *ξ* are constants. This subsampling leads to a bias: *m* (*η*^2^ Var[*At*]/ Var[*at*] − 1). Instead of where using time *t* and *t*+1, this method focuses on times *t* and *t*+*k* with different time lags *k =* 1, …, *k*_maximum_. With this, the branching ratio *m*_*k*_ is *< a*_*t*+*k*_ | *a*_*t*_ *>= m*_*k*_ *= η*^2^ Var [*A*_*t*_] / Var [*a*_*t*_] *m*^*k*^ *= bm*^*k*^, where *b* is a constant. To compute *m*_*k*_ with different *k*, we obtained an exponential curve and extracted *m* from this curve. *m* < 1 indicates a subcritical regime; *m* > 1 indicates a supercritical regime; *m =* 1 indicates a critical regime.

### 4.12 Convolution neural networks

#### Feedforward CNN

We used ResNet-18 (*36*) as FF-CNN. To calculate its eigenspectra, we used the pre-trained version on ImageNet provided by PyTorch. To evaluate its robustness against pixel noise, we trained ResNet-18 on MNIST with the Adadelta optimizer. The batch size was 64; learning rate was 1; weight decay was 0.0001; the coefficient used for computing a running average of squared gradients was 0.9; the term added to the denominator to improve numerical stability was 1 × 10^*−*6^, the number of training epochs was 10.

#### Recurrent CNN

We used the gated recurrent convolution neural network (*24*) as R-CNN, inspired by abundant recurrent connections in the visual systems of animals. The gates control the amount of context information inputted to the neurons. We used the code, GRCNN-55 (weight sharing), in https://github.com/Jianf-Wang/GRCNN. To calculate its eigenspectra, we used the pre-trained version on ImageNet provided by (*24*). To evaluate its robustness against pixel noise, we trained R-CNN on MNIST with the stochastic gradient descent (SGD) optimizer. The batch size was 64; the learning rate was 0.1; the momentum was 0.9; the weight decay was 0.0001; the number of training epochs was 10.

### 4.13 Eigenspectrum analysis

#### cvPCA

Eigenspectra of Billeh et al. model were estimated by the explained variance of the neural response along with the *nt*h principal component (computed from the first presentation). It is achieved by cvPCA that computes the covariance of the projections of neural responses for the two repeats onto this component. cvPCA measures the reliable variance of stimulus-related dimensions, excluding trial-to-trial variability from unrelated cognitive and/or behavioral variables or noise. It accomplishes this by computing the covariance of responses between two presentations of an identical stimulus ensemble (Fig. S2A). Because only stimulus-related activity will be correlated across presentations, cvPCA provides an unbiased estimate of the stimulus-related variance. Briefly, the algorithm operates as follows:

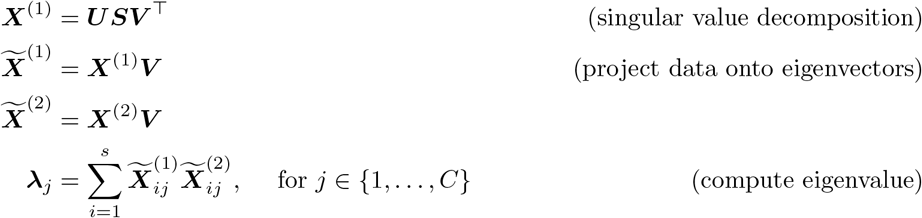

where ***X***^(1)^, ***X***^(2)^ ∈ ℝ^*S×N*^ (*S* is the number of stimuli and *N* is the number of neurons) are the neural responses for the first and second half of the trials (and averaged across trials), ***V*** ∈ ℝ^*N × C*^ are the *C* eigenvectors of the covariance matrix of ***X***^(1)^ and ***λ*** ∈ ℝ^*C*^ are the cross-validated eigenvalues associated with each of the eigenvectors (***λ***_***j***_ is the ***j*** th eigenvalue).

The first step of the cvPCA algorithm computes the eigenvectors of the neural response covariance from one set of the trials. The second and third steps project the neural responses from each half of the trials onto each eigenvector. The final step computes the (scaled) variance of the neural responses when projected onto an eigenvector (that was computed using one half of the trials). Thus, each cross-validated eigenvalue is related to the amount of stimulus-related variance of the neural responses along the eigenvalue’s corresponding eigenvector.

To be consistent with (*11*), we summed up spikes over 500 ms in response to visual stimuli. We ran cvPCA ten times on the response of the neural network fed with the same images that are used in (*11*). On each iteration randomly sampling the population responses of each stimulus from the two repeats without replacement. We ran ten different runs and found they were very similar to each other, i.e, the SD was close to 0. For the trained Billeh et al. model, we calculated the eigenspectra in three models trained with different noise and randomly generated data, and found the SD is 5.95 × 10^*−*5^. The displayed eigenspectra of the trained Billeh et al. model were averaged over these three models. The code is available in https://github.com/MouseLand/stringer-pachitariu-et-al-2018b.

#### Billeh et al. model

We analyzed the neural representation in the trained Billeh et al. model in the same way as responses of V1 neurons were analyzed in (*11*): Without loss of generality, we used 2,800 generic images which were randomly drawn from ImageNet validation dataset in all panels of Fig. 4. We also tried the 2,800 nature images used in (*11*) and found they gave rise to a slower decaying speed of eigenspectrum (1.15); there were only used in Fig. S2B to compare with the noise level in mouse V1 experiment. We also used a smaller set of 32 images, repeated 90 times. All stimuli were input 50 ms after the simulation onset and sustained for 500 ms in each trial to be the same as experimental procedures. They were presented twice to allow cross-validated analysis. The initial condition of membrane potentials and spikes was set to zeros, unless otherwise stated. We input the 2,800-nature-image stimuli 5 times with different random seeds that were used to draw the noise and initial conditions of membrane potential and after-spike current from uniform distributions. We found that the results were not sensitive to the initial condition and noise.

#### CNNs

Generic images were resized so that their shorter dimension was 256 pixels and then center-cropped to 224 × 224 pixels. Padding the image and resizing it to 224 × 224 pixels achieved similar results. Images were additionally preprocessed by normalizing each image channel (RGB channels) using the mean and standard deviation that was used during model training (mean: 0.485, 0.456, 0.406 and SD: 0.229, 0.224, 0.225). For gray images, we repeated it to 3 channels. Using these preprocessed images, we extracted activation from every layer of each CNN and computed their eigenspectra using principal components analysis (PCA), because artificial neural responses are deterministic.

#### Power-law fitting of eigenspectral

Using the least-squares method, we fit power laws to the eigenspectra, *f*(*n*), against PC dimension, *n*. The fitting function is *f*(*n*) = *n*^*−α*^(*n* ∈ [*n*_min_, *n*_max_]), where *n*_min_ and *n*_max_ are lower and higher bounds, respectively. For most cases, we chose *n*_min_ ∈ [1, 20] and *n*_max_ ∈ [301, 2800]. For the 32-grating recordings, owing to noise and the length of the spectrum, we chose *n*_min_ ∈ [1, 10] and *n*_max_ ∈ [14, 35]. For each possible pair of *n*_min_ and *n*_max_, we estimated the exponent *α* and its goodness-of-fit by the coefficient of determination (*R*^2^). We then selected as our estimate of *n*_min_, *n*_max_, and *α* that gave the maximum *R*^2^ (> 0.99) over all possibilities.

### 4.14 Discriminability index *d′* for neural responses to visual stimuli

To estimate how much information the neural activity conveyed about the stimulus identity, following (*3*), we used the metric *d*′, which characterizes how readily the distributions of the neural responses to the two different sensory stimuli can be distinguished (*37*). The quantity (*d*′)^2^ is the discrete analog of Fisher information (*38*).

**Fig. 6A and E**. To be consistent with the experimental study (*3*), we calculated the neural response as the spike counts in each bin of 200 ms and evaluated two different approaches to compute *d*′ values for the discrimination of the two different visual stimuli (gratings in the fine orientation discrimination task); the difference between two gratings is 2 *°*; each stimulus was presented in 500 trials. We analyzed the neural responses in a specific time bin relative to the onset of visual stimulation, which was called as instantaneous decoding approach used in (*3*). The alternative way, cumulative decoding, i.e., analyzing neural responses that were concatenated over time from the start of the trial up to a chosen time, demonstrated similar results. To determine *d*′ accurately despite having about fewer trials than neuron number in the Billeh et al. model, we reduced dimensional by using partial least squares (PLS) analysis (*39*) to identify and retain only 5 population vector dimensions in which the stimuli were highly distinguishable as in (*3*). In this 5-dimensional representation, the neural dynamics evoked by the two stimuli become distinguishable over the first 200 ms of stimulus presentation. In the reduced space, we calculated the (*d*′)^2^ value of the optimal linear discrimination strategy as:

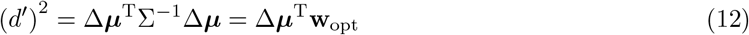

where 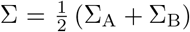 the noise covariance matrix averaged across two stimulation conditions, Δ***µ*** = ***µ***_A_ ***µ***_B_ is the vector difference between the mean ensemble neural responses to the two stimuli and **w**_opt_ = Σ^−1^Δ***µ***, which is normal to the optimal linear discrimination hyperplane in the response space (*38*). Each entry of a covariance matrix is the covariance of spike counts of two neurons *a*_*i*_ *a*nd *b*_*i*_ (*i* ∈ {1, 2, …, *N*}, *N* is the number of trials): 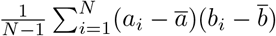 where 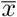 is the mean of {*x*_*i*_}.

We also calculated 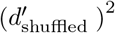, the optimal linear discrimination performance using trial-shuffled datasets, which we created by shuffling the responses of each cell across stimulation trials of the same stimulus.

Owing to this shuffling procedure, the off-diagonal elements of Σ_A_ and Σ_B_ became near zero. 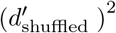 increased much faster with the increase of sampled neurons than (*d′*)^2^.

**Fig. 6B**. Noise eigenvalue *λβ* and its eigenvector 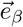 was calculated by eigen-decomposition of noise covariance matrix Σ.

**Fig. 6C**. To quantify the signals projected onto the eigenvector 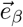, we projected Δ***µ*** onto 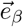 and calculated its norm |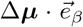|

**Fig. 6D**. To demonstrate that training makes signaling dimensions more orthogonal to the largest noise dimension, we decompose (*d*′)^2^ into a sum of projections:

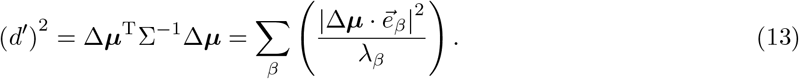

Because 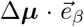 corresponds to the signal projected on noise eigenvector 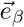 and noise eigenvalue *λβ* corresponds to the noise scale on 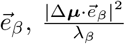 can be interpreted as singal-to-noise ratio on eigenvector 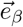. Clearly, the eigenvectors well aligned with Δ***µ*** are the most important for discriminating between the two distributions of neural responses.

## Acknowledgements

We would like to thank Eben Kadile, Du Kai, Oleg Kolner, Christoph Stöckl, and Yujie Wu for helpful comments on an earlier version of the manuscript. Also, we would like to thank Sandra Diaz for advice and help regarding large-scale computations. Most of computations were carried out on the Human Brain Project PCP Pilot Systems at the Jülich Supercomputing Centre, which received co-funding from the European Union (Grant Agreement number 604102). We also would like to thank the Beijing Academy of Artificial Intelligence for providing computational resources. This research was partially supported by the Human Brain Project (Grant Agreement number 785907) of the European Union and a grant from Intel.

## Supplementary Information for

**Table S1:** Performances on 5 tasks are not sensitive to the noise level. The light cyan highlights the noise level used in this study unless otherwise stated. VCDN is tested on novel images. FOD, fine orientation discrimination; IC, image classification; VCDN, visual change detection of nature images; VCDG, visual change detection of gratings; EA, evidence accumulation.

|  | FOD | IC | VCDN | VCDG | EA |
| --- | --- | --- | --- | --- | --- |
|  | acc. | acc. | acc. | acc. | acc. |
| $q = 1, s = 0$ | 93.67% | 89.14% | 83.61% | 92.37% | 93.75% |
| $q = 1, s = 2$ | 93.09% | 88.49% | 83.42% | 89.22% | 92.92% |
| $q = 1, s = 3$ | 93.47% | 88.97% | 82.52% | 88.73% | 90.17% |
| $q = 1, s = 3.5$ | 93.73% | 88.49% | 81.11% | 88.58% | 91.00% |
| $q = 1, s = 4$ | 93.06% | 88.81% | 82.30% | 87.86% | 91.50% |
| $q = 2, s = 2$ | 93.15% | 88.92% | 83.25% | 89.25% | 90.92% |
| $q = 2, s = 3.5$ | 93.93% | 89.19% | 82.20% | 87.51% | 90.58% |
| $q = 3, s = 3$ | 94.10% | 88.81% | 81.87% | 89.52% | 92.25% |
| $q = 3, s = 4$ | 93.53% | 89.02% | 83.06% | 88.14% | 89.33% |
| $q = 4, s = 3$ | 93.91% | 88.91% | 82.53% | 88.16% | 91.33% |
| $q = 4, s = 4$ | 93.47% | 88.16% | 82.16% | 88.17% | 89.08% |
| $q = 5, s = 4$ | 93.32% | 88.49% | 81.41% | 89.12% | 90.58% |
| $q = 10, s = 10$ | 94.03% | 88.54% | 82.97% | 89.56% | 84.33% |
| $q = 20, s = 20$ | 91.01% | 86.63% | 77.41% | 85.95% | 77.75% |
| $q = 200, s = 200$ | 54.58% | 39.56% | 53.09% | 53.96% | 52.67% |

**Figure S1:**
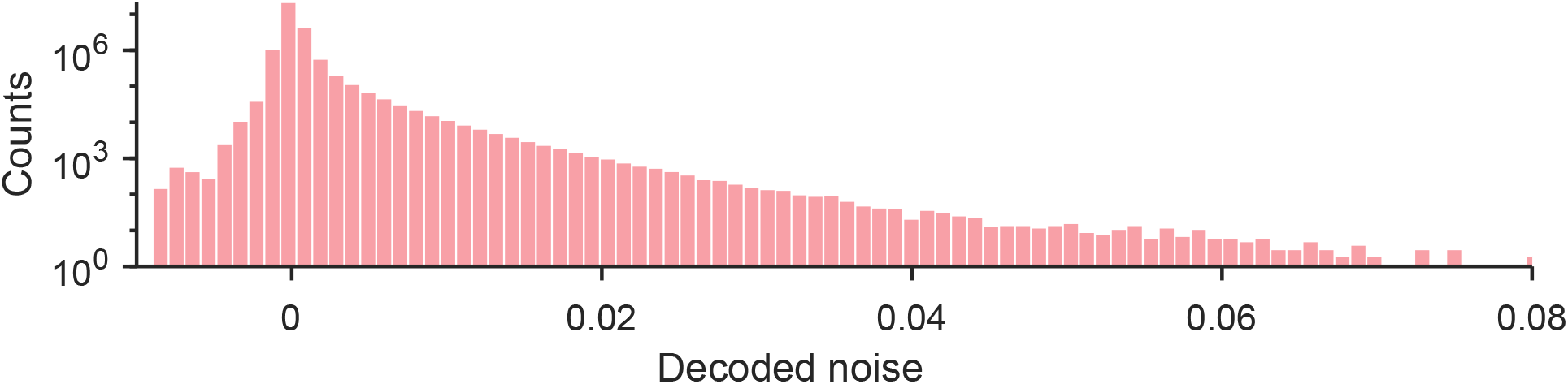
Empirical noise distribution where we drew the noise values. They are based on experimental data from (*11*).

**Figure S2:**
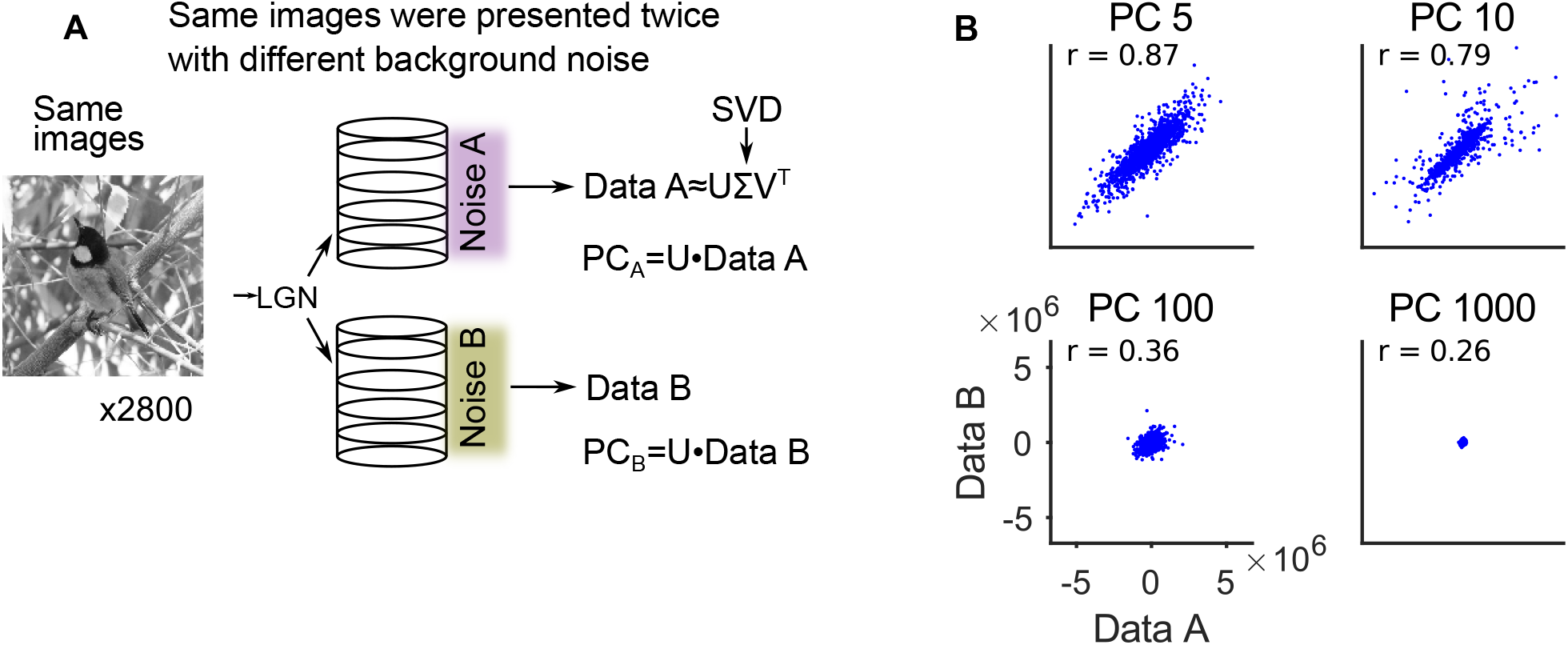
Schematic diagram of cvPCA (cross-validated PCA) to extract signal and noise. **(A)** Schematic diagram of cvPCA (cross-validated PCA). A set of generic images was presented twice with independently drawn noise for each image (Noise A and Noise B). Neural responses to the first presentation (Data A) were factorized by singular value decomposition (SVD) to estimate eigenspectrum of neural responses. **(B)** Correlations (r) of neural responses for two presentations of the same natural image, using our data-based noise model with *s = q =* 2 and projected onto selected principal components (PC).

**Figure S3:**
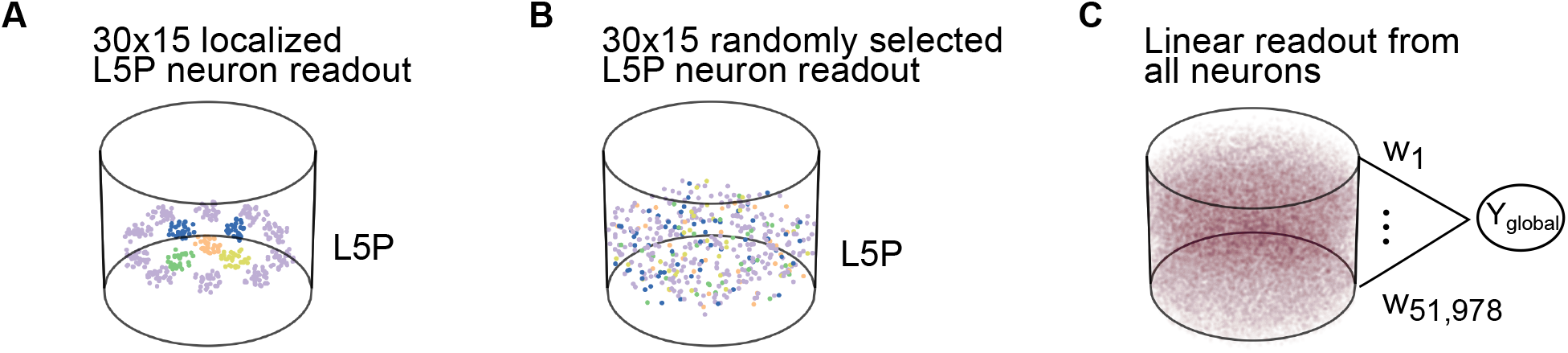
Schematic diagram of difference readout scenarios. **(A)** 15 spatially separated groups of 30 pyramidal neurons in L5 were selected to signal specific network outputs for 5 different tasks. Color encodes for 5 chosen tasks, same as in Fig. 2A. **(B)** Alternative selection of these 15 populations in L5 without spatial clustering leads to very similar performance. **(C)** Schematic diagram of a linear readout receives synaptic input from all 51,978 neurons in the microcircuit model, using a corresponding number of weights that can all be optimized for one particular task.

**Figure S4:**
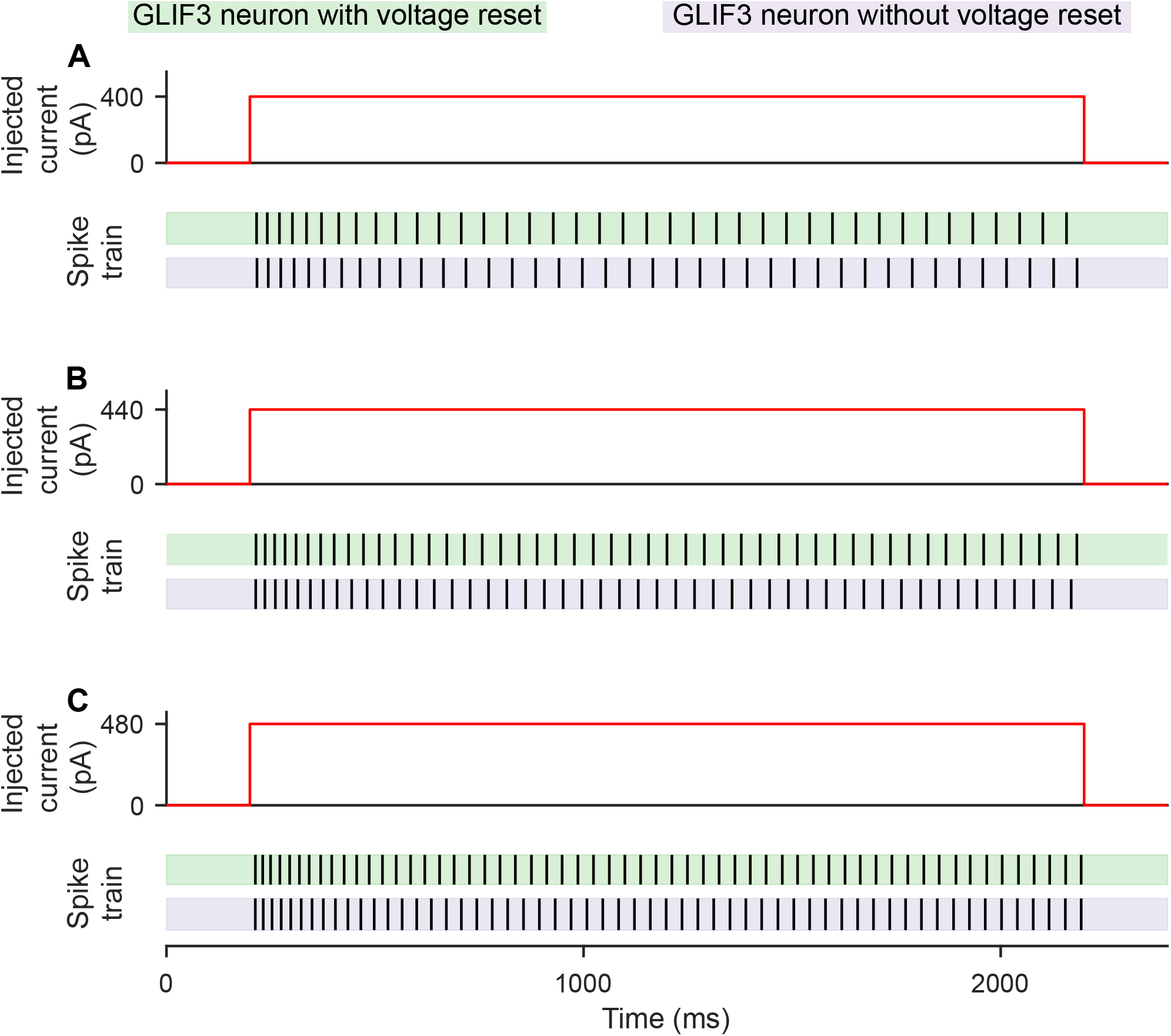
Adding voltage rest does not significantly affect the GLIF_3_ model. **(A)** Stimulus in form of a step function (top) was inputted to the GLIF_3_ model with voltage reset used in this study (middle) and the GLIF_3_ model without voltage reset in (*28*). **(B-C)** Same as in **(A)** but for two different intensities of step stimuli. The difference between two models is so small that it can be ignored. Henceforth, we call our modified GLIF_3_ model with voltage reset also as GLIF_3_ model for convenience.

**Figure S5:**
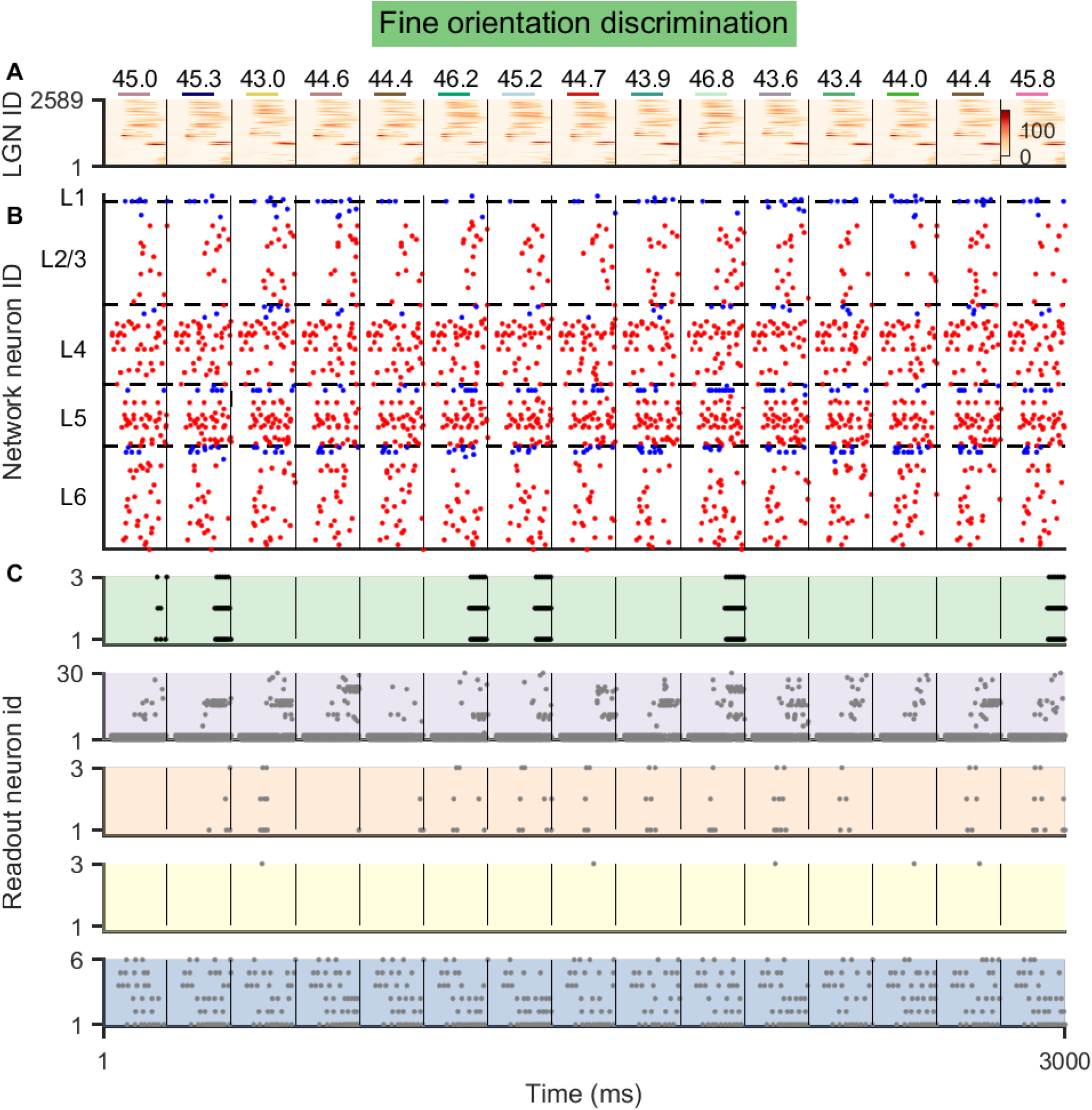
After learning, the model reliably distinguishes the orientation of gratings in the fine orientation discrimination task. **(A)** Colorful lines represent the timing of input images. Numbers on them represent the orientations of input gratings. The bottom colormap demonstrates the activity of LGN neuron activity. **(B)** Spike raster of the laminar V1 model. 200 neurons are sampled. Red and blue dots represent the spikes of excitatory and inhibitory neurons, respectively. Note that the spike and membrane potential of the model was reset to 0 after one classification was done (separated by the think black line). **(C)** Spike raster of readout neurons. 10% of neurons are sampled in every readout population. Color codes of panels are the same as in Fig. 2A. From the top to bottom, there are readout populations of the fine orientation discrimination, the image classification, the visual change detection of nature images and gratings, and the evidence accumulation tasks.

**Figure S6:**
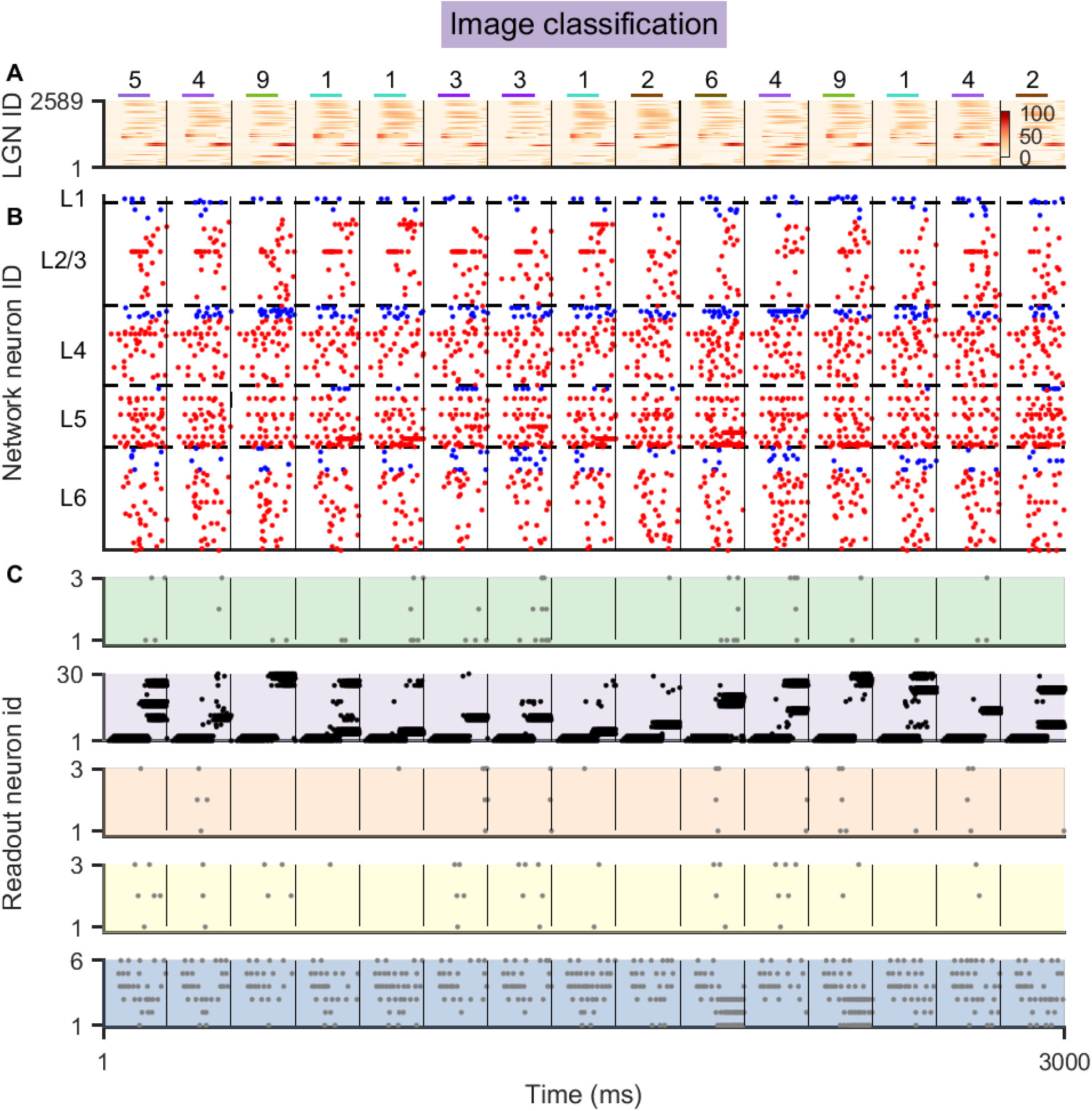
After learning, the model reliably classified the MNIST images. **(A)** Colorful lines represent the timing of input images. Numbers on them represent the digits in the input images. The bottom colormap demonstrates the activity of LGN neuron activity. **(B)** Spike raster of the laminar V1 model. 200 neurons are sampled. Red and blue dots represent the spikes of excitatory and inhibitory neurons, respectively. Red and blue dots represent the spikes of excitatory and inhibitory neurons, respectively. Note that the spike and membrane potential of the model was reset to 0 after one classification was done (separated by the think black line). **(C)** Spike raster of readout neurons. 10% of neurons are sampled in every readout population. Color codes of panels are the same as in Fig. 2A. From the top to bottom, there are readout populations of the fine orientation discrimination, the image classification, the visual change detection of nature images and gratings, and the evidence accumulation tasks.

**Figure S7:**
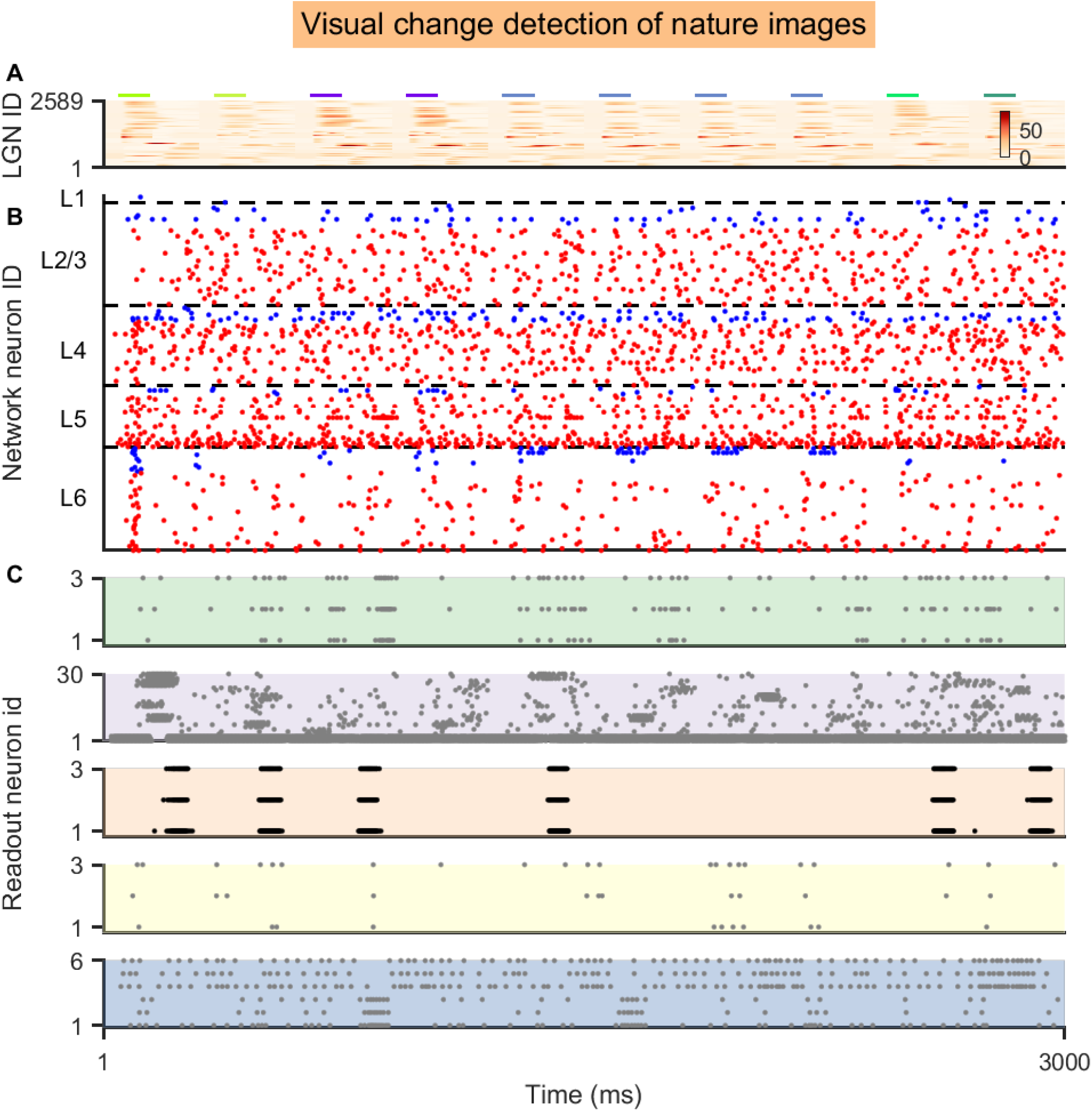
After learning, the model reliably reported the image identity change in the visual change detection task. **(A)** Colorful lines represent the timing of input images and the colors code the image identity. The bottom colormap demonstrates the activity of LGN neuron activity. **(B)** Spike raster of the laminar V1 model. 200 neurons are sampled. The slow noise was resampled every 600 ms. Red and blue dots represent the spikes of excitatory and inhibitory neurons, respectively. **(C)** Spike raster of readout neurons. 10% of neurons are sampled in every readout population. Color codes of panels are the same as in Fig. 2A. From the top to bottom, there are readout populations of the fine orientation discrimination, the image classification, the visual change detection of nature images and gratings, and the evidence accumulation tasks.

**Figure S8:**
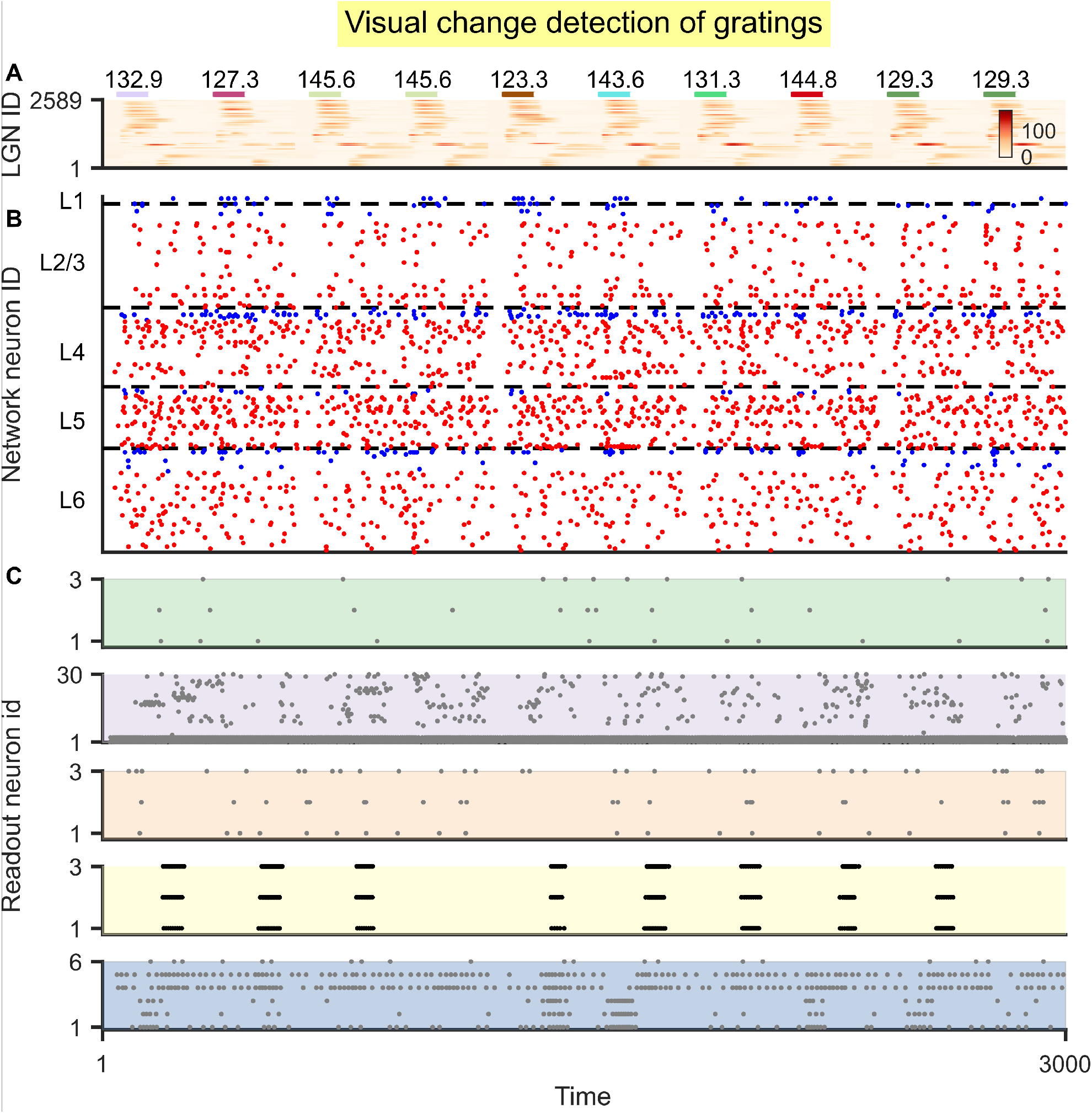
After learning, the model reliably reported the image identity change in the visual change detection task. **(A)** Colorful lines represent the timing of input images and the colors code the image identity. Numbers on them represent the orientations of input gratings. The bottom colormap demonstrates the activity of LGN neuron activity. **(B)** Spike raster of the laminar V1 model. 200 neurons are sampled. The slow noise was resampled every 600 ms. Red and blue dots represent the spikes of excitatory and inhibitory neurons, respectively. **(C)** Spike raster of readout neurons. 10% of neurons are sampled in every readout population. Color codes of panels are the same as in Fig. 2A. From the top to bottom, there are readout populations of the fine orientation discrimination, the image classification, the visual change detection of nature images and gratings, and the evidence accumulation tasks.

**Figure S9:**
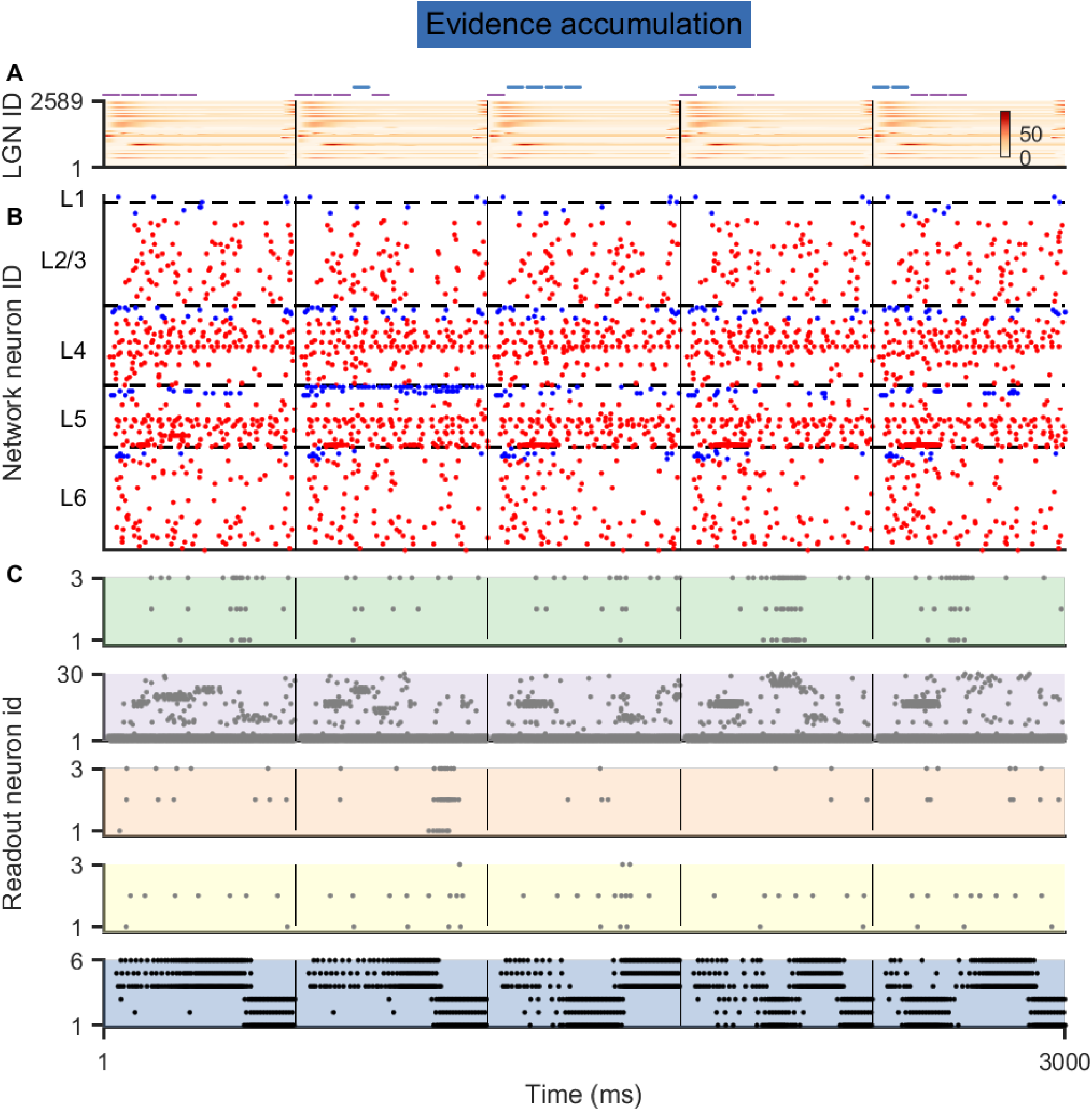
After learning, the spiking activity in the evidence accumulation task. **(A)** Colorful lines represent the timing of input left/right cues. The bottom colormap demonstrates the activity of LGN neuron activity. **(B)** Spike raster of the laminar V1 model. Red and blue dots represent the spikes of excitatory and inhibitory neurons, respectively. Note that the spike and membrane potential of the model was reset to 0 after one classification was done (separated by the think black line). **(C)** Spike raster of readout neurons. 10% of neurons are sampled in every readout population. Color codes of panels are the same as in Fig. 2A. From the top to bottom, there are readout populations of the fine orientation discrimination, the image classification, the visual change detection of nature images and gratings, and the evidence accumulation tasks.

**Figure S10:**
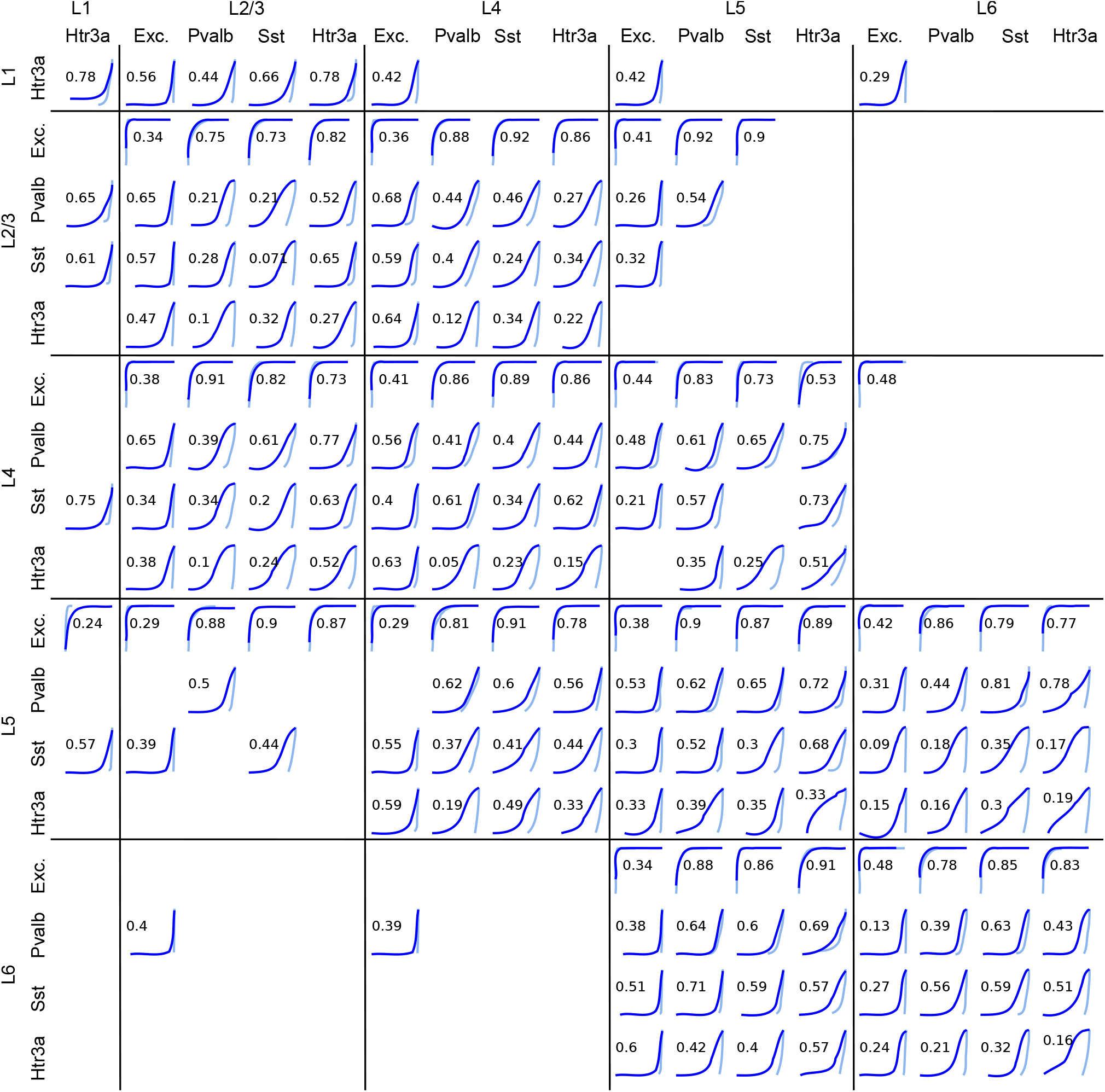
Distribution of recurrent weights between each population before (light blue) and after learning (dark blue) the task. Each row represents a pre-synaptic neuron population, and each column represents a post-synaptic neuron population. The histogram represents the distribution of synaptic weights of all synaptic connections that share the same pre-synaptic and post-synaptic neuron population. Vertical axis in each panel is log-scale. Horizontal axis is linear scale and horizontal range is from the smallest value to the largest value of each population. The number is 1 − *D* where *D* is from the Kolmogorov–Smirnov test, quantifying the similarity between distributions (*1*). Exc., excitatory neurons.

### 4.15 Supplementary Note 1

#### The differences of readout scenarios can explain why behavioral performance lags behind neural coding fidelity in area V1

The behavioral discrimination threshold for orientations in the mouse V1 was almost 100 times larger than the discrimination threshold which they inferred from neural coding fidelity of populations of 50,000 neurons in area V1 of the mouse (*4*). They conjectured that this difference was caused by the limitations of downstream decoders. The Billeh et al. model suggests a slightly more refined explanation. Direct measurements of coding fidelity based on simultaneous recordings from 50,000 neurons do not account for the fact that their information content has to be extracted by neurons in V1 that project to downstream areas. They are conceptually similar to the postulate of having a global readout neuron that receives synaptic input from all 50,000 neurons, see Fig. S3C. However, one can demonstrate in the Billeh et al. model that such a global linear readout attains for the fine orientation discrimination task an accuracy of 98.81%. On the other hand, a pool of 30 projection neurons on L5 could only achieve an accuracy of 93.15% if one assumed that they were localized closely together (Fig. S3A), and of 93.61% if they were assumed to be randomly distributed in L5 (Fig. S3D). These results suggest that how information from area V1 is extracted and projected to downstream areas is a limiting factor that is likely to contribute to the gap between the performance of an ideal observer of neural activity in V1 and the behavioral performance of mice.

### 4.16 Supplementary Note 2

#### Biological features speed up gradient descent training of spiking neural networks

We applied the same training procedure also to control models that lacked salient structural features of the Billeh et al. model (Materials and Methods), and found that their task performance advances substantially slower. We found that during the training time, the Billeh et al. model achieved an approximately saturating task performance, whereas these control models (all models contain 51,978 neurons) were only able to reach a substantially lower task performance level in 6 epochs, see Fig. 3F. Substantially larger computing resources will be needed to determine the performance levels that these control models can eventually reach after sufficiently long training.

It had already been shown that neuron models with slower changing internal variables tend to enhance BPTT training, see Fig. 2D of (*27*), and Fig. 3C and Supplementary Movie of (*40*). Fig. 3C of the latter reference also shows that a similar training advantage holds for a biologically more plausible variant of gradient descent learning. But Fig. 3F shows that also the laminar connectivity structure of the Billeh et al. model contributes to its learning speed, even if the neuron models remain unchanged (yellow curve). One possible explanation is that the laminar structure enforces topographic maps between different layers, and hence tends to keep information more local within the network. The data-based rapid spatial decay of connection probabilities within a layer has a similar effect. This locality of information processing may facilitate learning through local learning mechanisms in such network. In contrast, in a generic randomly connected network without this connectivity structure, all information is continuously dissipated throughout the network, which is likely to impede the localization of processing errors and their correction.

It will be interesting to see whether this effect also arises for biologically more realistic learning methods, for example, *e-prop* (*40*), which we did not perform because training with such methods tends to take substantially more trials, and therefore exceeded our computing resources. But since we now know that the Billeh et al. model is able to carry out the 5 visual processing tasks that we considered, one can now look for various biologically more realistic ways in the network initialization and/or learning algorithm -also for smaller instances of the Billeh et al. model- that can induce a similar computing capability or neural coding features.

## Notes

### Competing Interest Statement

The authors have declared no competing interest.

### Summary of Updates

Updated version to suffice for the requirement of journal submission.

